# Extracting multiple surfaces from 3D microscopy images in complex biological tissues with the Zellige software tool

**DOI:** 10.1101/2022.04.05.485876

**Authors:** Céline Trébeau, Jacques Boutet de Monvel, Gizem Altay, Jean-Yves Tinevez, Raphaël Etournay

## Abstract

Efficient tools allowing the extraction of 2D surfaces from 3D-microscopy data are essential for studies aiming to decipher the complex cellular choreography through which epithelium morphogenesis takes place during development. Most existing methods allow for the extraction of a single and smooth manifold of sufficiently high signal intensity and contrast, and usually fail when the surface of interest has a rough topography or when its localization is hampered by other surrounding structures of higher contrast. Multiple surface segmentation entails laborious manual annotations of the various surfaces separately. As automating this task is critical in studies involving tissue-tissue or tissue-matrix interaction, we developed the Zellige software, which allows the extraction of a non-prescribed number of surfaces of varying inclination, contrast, and texture from a 3D image. The tool requires the adjustment of a small set of control parameters, for which we provide an intuitive interface implemented as a Fiji plugin. As a proof of principle of the versatility of Zellige, we demonstrate its performance and robustness on synthetic images and on four different types of biological samples, covering a wide range of biological contexts.

## Introduction

The interplay between gene regulatory networks and physical forces in driving collective cell behaviors is key to tissue morphogenesis during development and to tissue homeostasis throughout life. Recent quantitative studies of epithelial morphogenesis have begun to unravel the basic cellular and physical principles of tissue development, by providing the tools to integrate multiple scales of tissue dynamics [1–4]. These tools are instrumental to quantify how cell shape changes, cell divisions, cell rearrangements and cell extrusions contribute to tissue remodeling, and to establish data-driven computational models of tissue morphogenesis.

Quantitative analysis of an epithelium starts with the extraction of its apical surface from 3D-microscopy images (z-stacks of xy-optical sections) encompassing the volume immediately surrounding the epithelium. However, this is a difficult task because this surface is usually not flat (it is best modelled as a curved surface, or 2D submanifold embedded in 3D space), and is often surrounded by other biological structures such as cell layers, acellular membranes, extracellular matrix, and vesicles that hamper its visualization and reconstruction. To make the surface extraction tractable, these studies rely on specific preparations of the specimen, allowing to expose the entire epithelial surface labeled with junctional fluorescent markers to reveal the network formed by epithelial cell-cell contacts. Once the epithelial surface has been extracted, automated cell segmentation and cell contour tracking tools can be used to follow the dynamics of every cell within the epithelium.

Another challenging experimental limitation of these studies is that some of the structures surrounding the epithelium exert external physical constraints that are known to critically affect epithelial morphogenesis by directing cellular dynamics and signaling pathways [1, 5–7]. To understand the physical forces controlling tissue morphogenesis, it is thus essential to also characterize how the dynamics of these extra-epithelial surfaces relate to that of the epithelium (see [8] for review). This calls for the development of dedicated tools allowing the automated extraction of information from several surfaces of interest in a given sample, since the sheer volume of the data precludes any attempt at a manual analysis.

Several surface extraction tools have been developed, some of which are available as open access software, such as PreMosa [9], FastSME [10], and LocalZProjector [11]. These tools focus on the extraction of a single, near-horizontal epithelial layer, which is assumed to (i) be sufficiently smooth, (ii) show enough contrast against surrounding background signals, and (iii) cover the entire image field-of-view. Specifically, it is assumed that the fluorescent marker used to label the epithelial cell network should provide the highest contrast in the image and allow to select it out from autofluorescent extracellular structures such as the cuticle in flies, or other acellular membranes in mammalian epithelia. The surface is then localized using heuristic algorithms based on the detection of the pixels of maximum contrast and/or brightness. However, applying these tools on more complex biological images with several epithelia of weaker contrast often leads to incorrect localization of the surface of interest, and its blending with the nearby unwanted biological structures.

MinCostZ on the other hand, is the only available open-source tool that allows the extraction of up to two surfaces from a 3D stack, and imposes explicit continuity constraints on the reconstructed surfaces. MinCostZ surface extraction relies on a previously developed formulation of the problem as a graph-cut optimization [12]. It is implemented as an ImageJ plugin [13], taking as control parameters, the number of surfaces to extract, the maximum slope and the range of distances allowed between the surfaces, as well as some user-defined cost function that should reflect the characteristics of the surfaces in term of signal intensity, contrast and texture. Despite its interest, this approach remains computationally costly and difficult to apply in practice due to the non-trivial choice of the cost function and the need to know beforehand the relative positions of the surfaces to be extracted.

Alternatively, one can segment the surfaces of interest by using supervised machine learning tools such as the software solutions Weka [14] or Ilastik [15], as proposed in the ImSAnE surface reconstruction framework [16]. A deep learning approach, using a network of the U-net type to segment the pixels belonging to a single surface of interest, has also recently been reported [17]. While promising as they can provide state of the art segmentations of epithelial surfaces in difficult imaging conditions, machine learning approaches require the prior manual annotation of a sufficiently large set of surfaces to generate suitable training sets. This process can be very time consuming, often necessitating several rounds of trials and errors to obtain satisfactory results, without guarantees to be generic, *i.e.,* to generalize to a wide range of datasets. So far, no solution to the multiple surface extraction problem has been proposed, which is satisfactory both in terms of genericity and ease of use.

However, such a tool is highly desirable for modern biology studies. Indeed, tissue organization in the context of developmental biology emerges from the interaction of several neighboring structures [8] through the interplay of molecular signals [18, 19], as well as electrical [20, 21], hydraulic [22, 23] and mechanical contact interactions [24, 25]. The importance of such interaction is exemplified in embryonic explanted tissue cultures that develop abnormally when separated from their neighboring structures [26]. Similarly, in the context of tissue engineering, stem-cell derived aggregates harbor various types of tissues surrounding the genuine organoid, and these tissues presumably influence organoid shape, fate and differentiation (see [27] for review). The ability to simultaneously study the dynamics of neighboring structures together with the structure of interest is therefore essential for an integrated understanding of tissue development, and for any attempt to harness tissue self-organization *in vitro.*

Here, we introduce Zellige, a tool based on a novel constructive approach that allows the automatic extraction of a non-prescribed number of surfaces from a 3D image. To do this, the user is only required to adjust a small set of intuitive control parameters, a task largely facilitated by a user-friendly interface implemented as a plugin for the open-source Fiji platform [28]. We tested the performance and robustness of Zellige for multiple surface extraction by applying it to synthetic images and 3D microscopy images from four different types of biological samples, containing multiple surfaces of interest of widely varying texture and contrast. These experiments demonstrate the ability of the approach to extract several (up to 4) surfaces of potentially very low contrast, selectively from other highly contrasted and complex structures, with a single set of reconstruction parameters. A sensitivity analysis also reveals a high robustness of Zellige against small variations of these parameters. This will make it a tool of choice in terms of versatility and ease of use for the investigation of biological surfaces.

## Results & discussion

### Proof of concept of multiple surface extraction on a synthetic image

The implementation of Zellige is summarized in **Figure 1** and in the *Methods* section (**Figure S1**). **Figure 2** shows the results produced by Zellige on a phantom 3D image [29] containing three distinct synthetic surfaces generated as described in Supplementary note 2. The three surfaces are extracted with little errors. We assessed the quality of the reconstruction by comparing each of the height-maps produced by Zellige to the corresponding ground truth (GT) height-map, which is exactly known in this case (**Figure 2A-C**, and Supplemental note 3). For the three surfaces, the reconstruction has subpixel accuracy over >99% of the GT pixels (**Figure 2D-E**), with a root mean square error (RMSE) of ≤ 0.6 in pixel units, showing that the surface localization is highly accurate. In addition, the coverage, which measures the proportion of the reconstructed surface relative to the GT, is near 100% for the three surfaces. To achieve these results, the control parameters of the two steps of the surface extraction were adjusted manually to some adequate reference values (see Supplementary Table S1) using the Zellige Fiji interface. Only the parameters controlling the pixel classification step (amplitude and Otsu threshold parameters *T_A_* and *T*_otsu_, minimal island size *S*_min_, and smoothing parameters *σ*_xy_ and *σ*_z_) did actually require a modest adjustment. The parameters of the surface assembly step (parameters *T*_OSE1_, *R*_1_, *C*_1_ and *T*_OSE2_, *R*_2_, *C*_2_ of the 1^st^ and 2^nd^ construction rounds, respectively) were set to their default reference values (see **Figure S2** and Supplemental note 4) and did not need to be adjusted.

**FIGURE 1.**
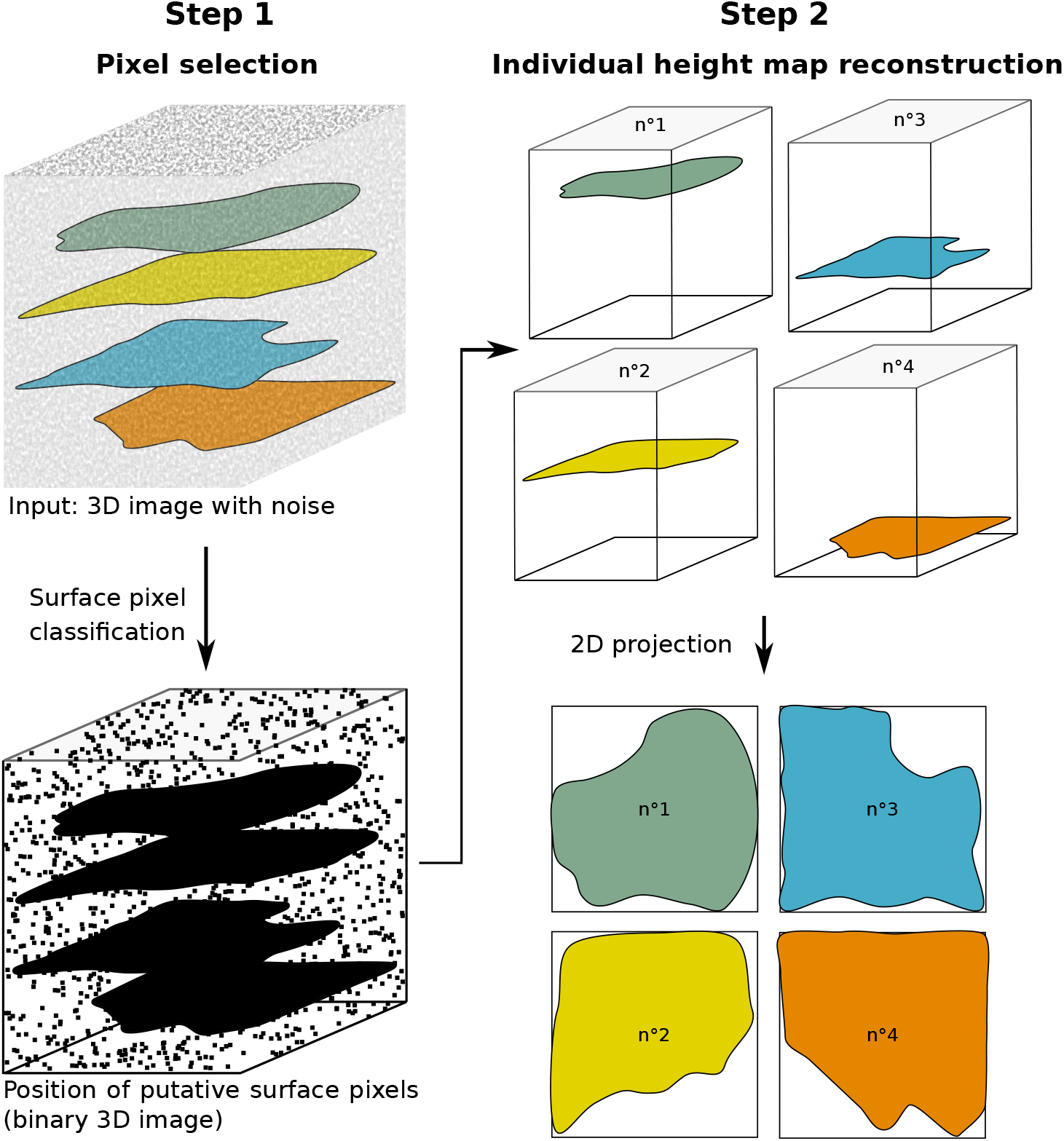
Flowchart of Zellige’s algorithmic steps. Surface pixel selection (step 1), surface assembly in the form of a height map (step 2), and subsequent projection localized to the height map, are schematically depicted in the case of a 3D image containing 4 surfaces of interest.

**FIGURE 2.**
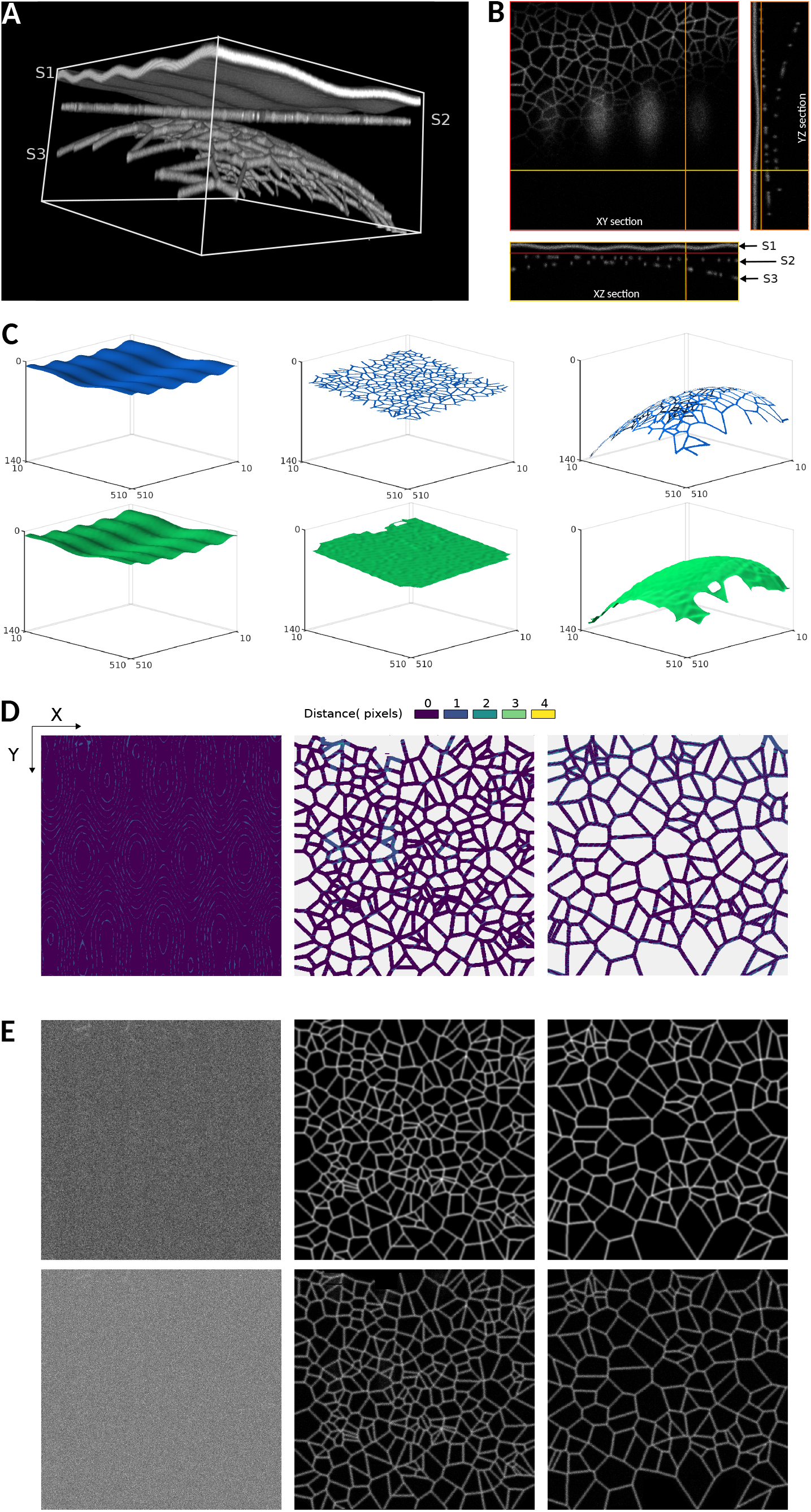
Multiple surface extraction on a synthetic 3D image. (A) The image contains 3 phantom surfaces (S1, S2, S3) of different shapes (sinusoidal, flat, and paraboloidal, respectively), and different textures (surface S1 has constant intensity, while surfaces S2 and S3 are supported by Voronoi meshes of different cell-sizes). (B) 3D representations of the height maps extracted by Zellige (in green) and of the ground truth (GT, in blue) height maps of surfaces S1, S2, and S3. (C) Error maps displaying the distance along the z-axis between the reconstructed and GT height maps for surfaces S1, S2, and S3. (E) Projections of the 3D image localized to the different surfaces S1-S3 (maximum intensity projections over a subvolume of a width δ z=1 pixel above or below the corresponding height-maps). Upper and lower panels show the projections based on the GT and the reconstructed height-maps, respectively.

Thus, using a single set of control parameters, Zellige can extract multiple surfaces of various shapes and textures with virtually no error, without requiring the user to provide information about their number or relative position, nor about their shape or texture characteristics.

### Performance of Zellige on biological samples

#### Example 1: Extracting multiples surfaces from an image of a pupal fly specimen

Over the past few decades, the *Drosophila* model has been invaluable to decipher the molecular and cellular mechanisms underlying organ embryogenesis [30, 31]. Epithelium morphogenesis studies not only revealed the importance of mechanical stresses (including stress boundary conditions) and planar polarity signaling on cell dynamics to generate tissues of reproducible sizes and shapes, it also highlighted the importance of extracellular matrix attachments in constraining the tissue stresses that guide patterning [1, 32]. At the pupal stage, the fly undergoes dramatic remodeling of its larval organs into adult organs. Large scale tissue flows initiate at a timing that coincides with molting, when the epithelium contracts away from the overlaying cuticular sac, a protective acellular membrane that imposes mechanical boundary conditions to the tissue.

**Figure 3** shows the results of applying Zellige on a 3D image of a *Drosophila* pupa acquired with a spinning disk confocal microscope [33]. The sample expresses Ecadherin-GFP, a fluorescent marker of cell-cell junctions, and encompasses a portion of the pupa’s abdomen and a small portion of its wing. Four surfaces of interest can be identified, with varying signal intensities, noise levels and features (**Figures 3A-B**). The abdomen is formed of an epithelium (surface S2) overlaid by a cuticle (surface S1). Lying just beneath these two surfaces, one can observe globular structures showing in some places a higher intensity than the signal coming from the surfaces. The wing also consists of an epithelium of low intensity signal (surface S4), and an overlying cuticle (surface S3). These two surfaces are relatively flat, except for surface S3 which is very steep near one of its edges.

**FIGURE 3.**
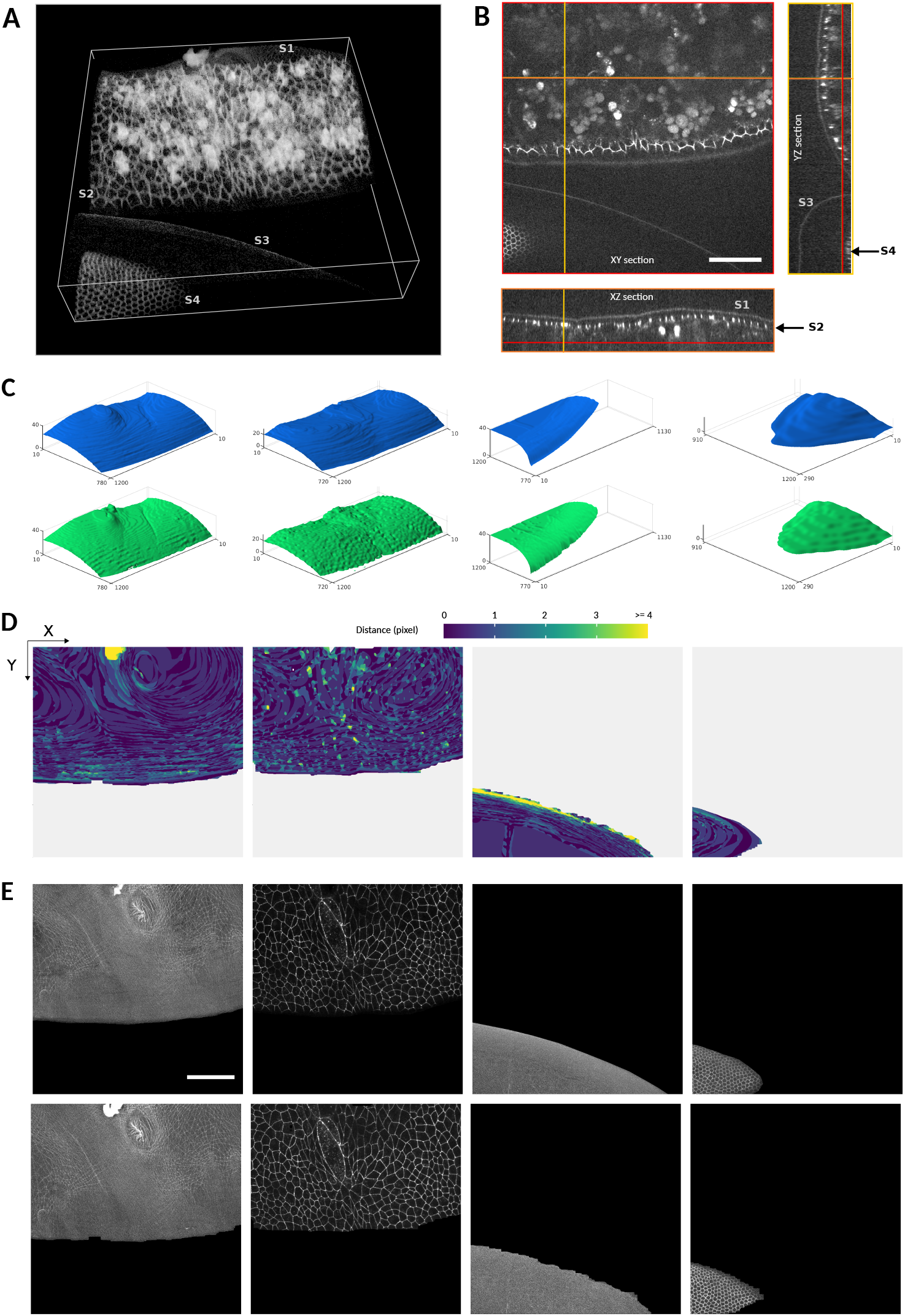
(A,B) Volume rendering (A) and orthogonal sections (B) of a 3D image of fly embryo taken around 24h after puparium formation, covering a portion of the abdomen (showing histoblast cells and larval cells), and a portion of the developing wing. Scale bar 50 μm. Four surfaces of interest may be identified in the dataset (of dimensions 1200 × 1200 × 51 pixels): surfaces S1 and S2 are relatively close to one another and located within overlapping z-ranges (8 ≤ z ≤ 50 and 20 ≤ z ≤ 50, respectively). Surfaces S3 and S4 (located in the z-ranges 42 ≤ z ≤ 50 and 9 ≤ z ≤ 50, respectively) are relatively far from each other and can nearly be separated by a plane. (C) 3D representations of the height maps extracted by Zellige (in green) and of the ground truth height maps (GT, in blue) of surfaces S1-S4. The reconstructed height-maps of all surfaces S1-S4 cover >93% of the area of the corresponding GT (cf. Figure S2 and **Table S1**). To reduce the staircase artifacts (more or less visible depending on the surface) due to the digitization of the GT and reconstructed height-maps, all height-maps were smoothed with a 2D gaussian filter with a standard radius of 5 pixels (cf. Supplemental note 1). (D) Error maps (color-coded distance along the z-axis between the reconstructed and the GT height-maps) plotted for each of the reconstructed surfaces. The large majority of pixels on the reconstructed height-maps (98%, 96%, 91%, and 99% for surfaces S1 to S4, respectively) display errors of <2 pixels. The height-maps of surfaces S1, S2, S4 show subpixel accuracy on average (RMSE < 1), while that of surface S3 is slightly less accurate (RMSE = 1.25). (E) Projections of the 3D image localized to the different surfaces S1-S4 (in this and all subsequent figures, these are maximum intensity projections over a subvolume of width δ z= ± 1 pixel above or below the corresponding height-maps). Upper and lower panels show the projections based on the GT and the reconstructed height-maps, respectively.

**Figure 3C** shows a 3D graphical representation of the height-maps reconstructed by Zellige (green) and those reconstructed by an expert biologist (blue), taken as ground truth (GT). While these height-maps clearly show greater roughness than those of the synthetic surfaces presented earlier, they could again be obtained with a single set of control parameters that were adjusted manually with the Zellige interface (see **Supplementary Table S1**). We observe an excellent match between the four reconstructed and corresponding GT height-maps, despite the rather complex topography of surfaces S1 and S2 (with slopes reaching up to 45°), and the near-vertical inclination of surface S3 at its boundary. Yet, small deviations may be seen in the regions of highest slope of the surfaces. Some of these deviations are likely attributable to uncertainties in the definition of the GT height-maps, whose accuracy depends on the expert.

**Figure 3D** shows the differences between the reconstructed and GT height-maps, plotted as color-coded error maps. These differences are <2 (in pixel units) for most pixels, while some regions of higher error values can be seen locally in surfaces S1 and S2, and at the boundary of surface S3. Note that for surfaces S2 and S4, which contain junctional epithelial meshes composed of larger and smaller cells, respectively, the GT height-map encompass not only the mesh but also the interior of the cells, where no junctional signal is detected. The distance calculated inside the cells is thus more subjected to intensity fluctuations, especially for surface S2. Nonetheless, the RMSEs of surfaces S1, S2 and S4 are less than 1, showing that on average the reconstructed height-maps match the corresponding ground truth with subpixel accuracy. The higher RMSE (1.25) of surface S3 is largely due to the region of steep region at the edge of this surface (yellow region on the error map for this surface, **Figure 3D**). The coverage of the reconstructed height-maps is excellent (≥ 96%) for surfaces S1 and S2, and slightly lower, but still very good (≥ 93%) for the smaller surfaces S3 and S4. **Figure 3E** shows the 2D projections of the 3D image obtained for each of the reconstructed surfaces and for the corresponding ground-truths. The inaccuracies visible on the error maps (see **Figure 3D**) do not significantly impact these projections, which appear very similar to the projections obtained with the corresponding ground-truths. Thus, while the biological sample contains significant noise and shows a much more variable contrast (especially with the presence of high intensity globular structures near surfaces S1 and S2), Zellige makes it possible to segment these surfaces selectively, with a quality of segmentation comparable to that obtained by manual expert segmentation.

This possibility brings several perspectives that are not offered by single surface extraction algorithms. First it opens the possibility to systematically study the tissue axial movements (along z) relative to the cuticle during molting, allowing for example to gain insights into the early tissue contraction of the wing hinge that acts as a mechanical inducer over the wing blade [8]. Second, Zellige makes it possible to automatically extract structures such as the abdomen epithelium, which is usually segmented manually [34], due to the difficulty to separate the large larval cells from the cuticle mesh and from other globular structures (such as fat bodies or macrophages) present underneath the epithelium. All these structures become intertwined when using other extracting tools. In this context, Zellige opens new opportunities to study collective cell behavior during epithelial morphogenesis *in vivo,* and to integrate in the analysis the surrounding surface-like structures involved in the mechanics of the system.

#### Example 2: Extracting a thin cochlear epithelium surface from a multilayer dataset

As the first model in which planar cell polarity signalling was shown to be conserved in vertebrates [35], the mammalian auditory organ, the cochlea, is arguably our most valuable model to study epithelial patterning and morphogenesis beyond the fly and zebrafish [36, 37]. Cochlear morphogenesis involves complex and tightly controlled patterning processes during which the cochlear sensory epithelium extends and develops its characteristic coiled snail shape, while adopting a striking cellular mosaic organization, with graded changes of morphogenetic parameters along the cochlea [38, 39]. These morphogenetic processes are well recapitulated in organotypic cultures, on the condition that the mesenchyme that underlies the epithelium be preserved. The cultures are then amenable to live imaging [37, 40], pharmacological [41] and genetic manipulations.

**Figure 4A** shows a confocal swept field microscope acquisition of an embryonic mouse cochlea [42]. The sample contains only one surface of interest, the cochlear epithelium, but this surface lays on top of a thick tangled mesh of non-epithelial cells originating from the mesenchyme. The whole biological tissue is stained for filamentous actin (F-actin) using phalloidin. The epithelium surface presents a non-uniform signal included in a small z-range (6 ≤ z ≤ 10), and a mesh of very heterogeneous size. Between sections z = 10 and z = 14 one can observe the basolateral region of the epithelial cells, also stained for F-actin. he particularity of this sample is that the mesenchyme presents an intense and contrasted signal over a wide range of z-values (14 ≤ z ≤ 43). This makes it challenging to extract the surface of the epithelium, which is characterized by low intensity and low contrast.

**FIGURE 4.**
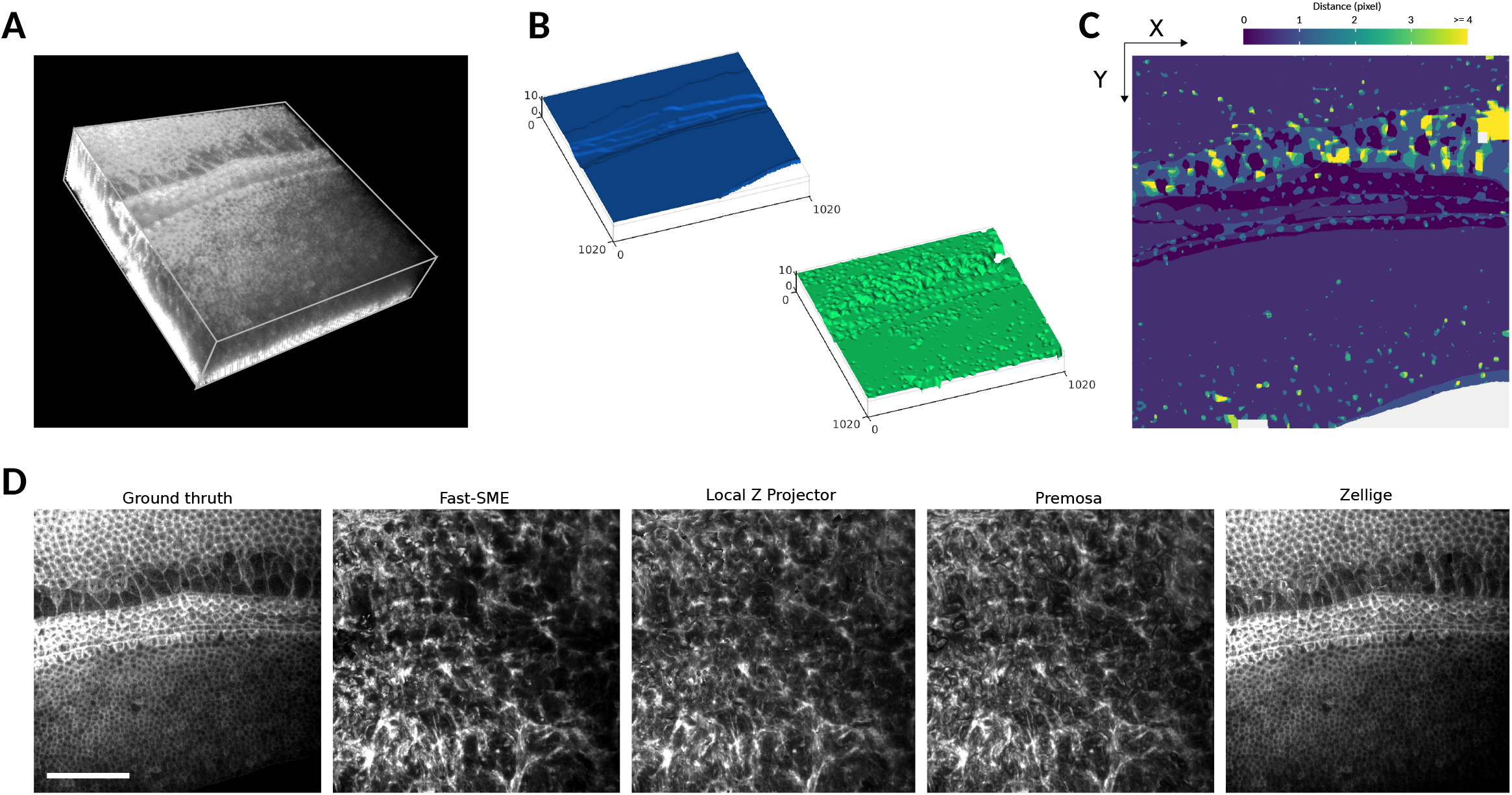
(A) Volume rendering of a 3D confocal swept-field image of the mouse cochlear embryo on embryonic day E14.5. The dataset (of dimensions 1024 × 1024 × 45 pixels) shows a portion of the sensory epithelium (at the topmost sections of the stack) and the underlying non-cellular layer of mesenchyme on which the organ develops. Both structures are stained with phalloidin to reveal F-actin. Scale bar 40 μm. The surface of interest is the epithelium surface, harboring the sensory and supporting cells under differentiation. The mesenchyme layer is not (strictly speaking) assimilable to a surface, but it produces a strong background signal nearby the surface of interest, hampering its extraction. (B) 3D representations of the height map extracted by Zellige (in green) and the GT height map (in blue), of the epithelium surface. (C) Color-coded error map of the reconstructed height-map, which shows subpixel accuracy (errors <1) over a large majority (83%) of pixels, as well as on average (RMSE ~ 1.1). (D) Projections localized to the GT height-map of the epithelium surface (left most panel), and to the height-maps extracted with the four different algorithms: FastSME, LocalZProjector, PreMosa, and Zellige. Only Zellige correctly extracts the surface of the epithelium in this example.

**Figure 4B** shows a 3D representation of the height-map reconstructed by Zellige and the corresponding GT height-map (again reconstructed manually by an expert). On the corresponding error map (**Figure 4C**), most (~83%) pixels of the reconstructed height-map show subpixel accuracy (with distances < 1 to the corresponding pixels of the GT height-map). The errors are greater in regions where the cell size is larger, as well as in the area where the signal intensity is very low. However, they remain smaller than 2 for > 95% of the pixels. This result is consistent with the low value of the RMSE (1.1). The surface is also reconstructed with an excellent coverage (> 99%, **Supplementary Table S1**).

As the sample contains a single epithelial surface of interest, we compared the performance of Zellige with three other software that can extract only a single surface (**Figure 4D**). The projections of PreMosa, FastSME and LocalZProjector completely miss the epithelium. Only regions of high contrast corresponding to the mesenchyme are projected. In contrast, Zellige generates a projection very close to the ground truth. This demonstrates the efficiency of Zellige to selectively extract a low contrast surface, despite the presence of several structures of higher contrast. Indeed, Zellige detects every structure as a possible surface seed without any assumption on its contrast, and only extends this seed into a surface if enough spatial continuity is found in the surrounding signal. This feature allows to separate individual surfaces from other structures spatially, which should greatly facilitate the analysis of live imaging experiments.

#### Example 3: Extracting a single bronchial epithelial surface rendered abnormally rough by SARS-CoV-2

Recently, we used Zellige to extract the surface of a primary culture of bronchial epithelial cells following infection by the SARS-CoV-2 virus [43]. The infection causes the surface of the epithelium to become abnormally rough due to cell damages as seen from discontinuities within the cell layer. The sample we chose from this study is a 3D confocal image of the epithelium responding to SARS-CoV-2 infection [44] (**Figure 5A**). The surface of interest in this image corresponds to the layer of epithelial cells stained for the tight junction protein Zona Occludens-1 (ZO-1). The surface roughness causes the network of junctions to extend over the height of the z-stack, with a signal of varying intensity (**Figure 5A**). In addition, the junctional network remains non-planar even at the level of a single cell, hence violating the smoothness condition commonly assumed to hold in the context of epithelial surface extraction. We also observe the presence of nearby punctiform structures of high contrast that are mainly located outside of the epithelium surface. This sample therefore provides an example of a surface with a complex landscape, interspersed with a constellation of signals which may interfere with the surface segmentation.

**FIGURE 5.**
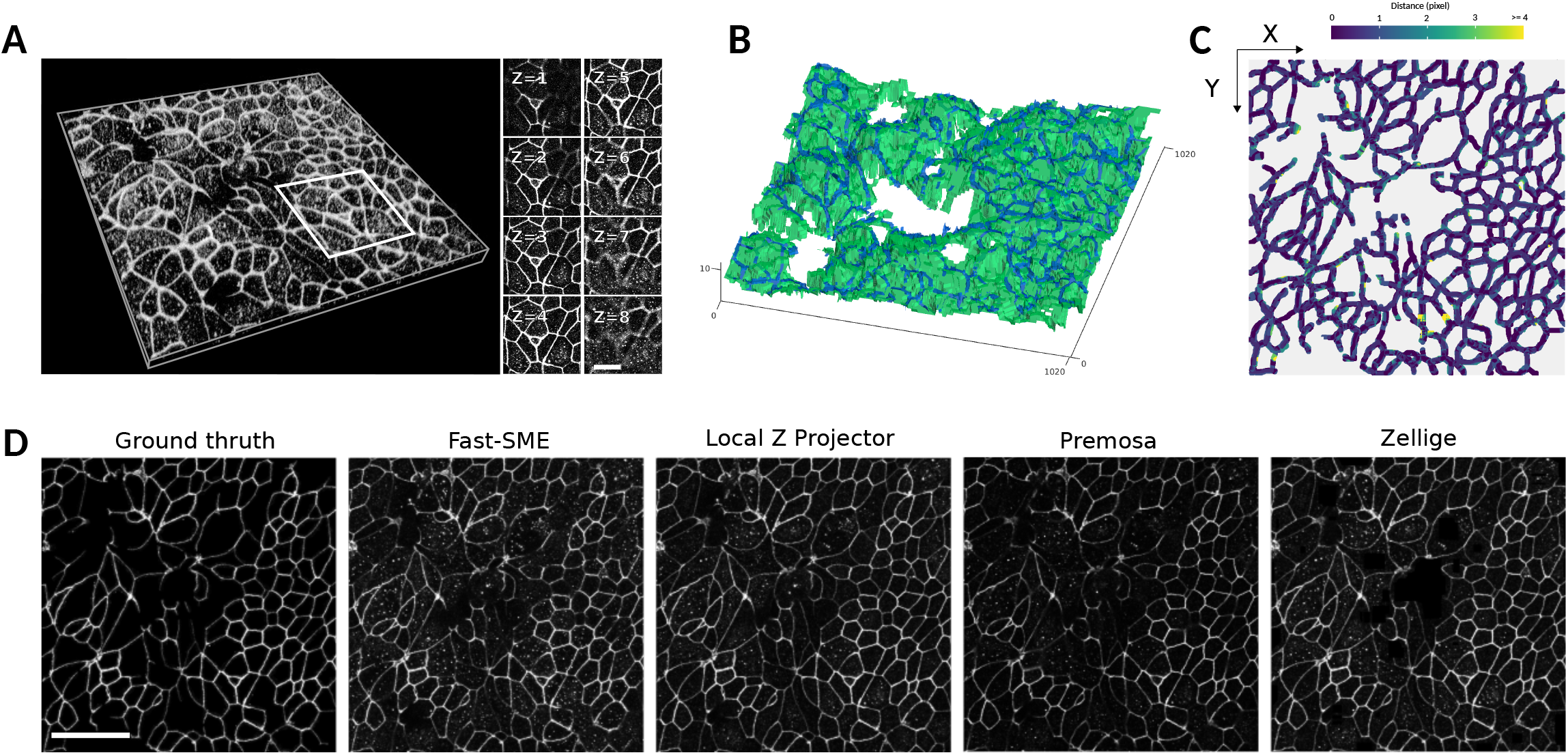
(A) Volume rendering and individual sections of a confocal 3D image of a primary culture of bronchial epithelial cells 4 days after it was infected by the SARS-CoV-2 virus. The dataset (of dimensions 1024 × 1024 × 15 pixels) covers a portion of the epithelium immunostained for the tight junction protein ZO-1. Notice the roughness of the epithelium surface and the presence of anomalous bulges (arrows) resulting from the SARS-CoV-2 infection. Scale bar 10 μm. (B) 3D representations of the height map extracted by Zellige (in green) and the GT height map (in blue), of the epithelium surface. (C) Color-coded error map of the reconstructed height-map. Despite its roughness, the surface of interest is reconstructed with subpixel accuracy over the majority (71%) of pixels, as well as on average (RMSE ~ 0.81). (D) Projections localized to the GT height-map of the epithelium surface (leftmost panel), and the height-maps extracted with the four different algorithms: FastSME, LocalZProjector, PreMosa, and Zellige. Scale bar 30 μm

The 3D representation of the reconstructed and corresponding GT height-maps (**Figure 5B)** makes it possible to appreciate the roughness of the surface of interest. The two height-maps overlap quite satisfactorily. As shown on the error map (**Figure 5C**), a large majority (71.1%) of pixels of the reconstructed height-map show errors smaller than 1 pixel (~ 96% of them showing errors smaller than 2 pixels). The error is however larger in regions where the cell size is larger, as well as in areas where the ZO-1 signal intensity is very low, preventing a complete reconstruction of the junctions. Nevertheless, the overall RMSE remains small, with a value of 0.81 (**Figure 5**). The coverage of the reconstructed surface is also excellent (98% of the GT height-map), despite the above-mentioned discontinuities. **Figure 5D** shows the comparison of the projection generated by Zellige to those produced by PreMosa, FastSME, and LocalZProjector. The projection generated by PreMosa misses many junctions of the epithelial network, but it removes quite well the punctiform signal originating from other optical sections. FastSME performs better than PreMosa in reconstructing the junctions, but they produce a projection where the punctiform signal remains strong. In contrast, Zellige and LocalZProjector manage to both reconstruct the surface well and to filter out the punctiform signal quite effectively. This result demonstrates the efficiency of Zellige to extract a surface with complex topography by excluding intense and contrasted spurious signals away from the epithelium surface.

#### Example 4: Extracting the apical and basal layers of a dome-shaped epithelium (developing inner ear organoid)

Organoids are stem cell-derived and self-organizing 3D tissue structures that can mimic certain organ structures. They have emerged as promising *in vitro* models for developmental biology research, as well as biomedical translational research applications. Here we take the example of mouse stem cell derived inner ear organoids that form vesicular structures composed of an epithelium harboring sensory cells. These organoids are part of a cellular aggregate that also contains other tissues such as the mesenchyme [45] adjacent to the organoids. The epithelial cells of the forming inner ear organoids acquire a basal-apical polarity, with their apical side facing the lumen of the organoid, and their basal side facing outwards. The apical junctional network of the epithelium is difficult to visualize in microscopy images as it is seen from below, through the basal layer. Another difficulty is the spherical geometry of the vesicle system, which makes the epithelial surface of interest difficult to extract in regions of high inclination relative to the focal plane.

**Figure 6** shows the result of applying Zellige on a 3D confocal microscopy image of half of a developing inner ear organoid at 14 days of culture, a stage at which markers characteristic of the mouse otic vesicle can be detected [46]. The sample was fixed and stained for F-actin to visualize all cellular structures including the epithelium. Two surfaces of interest can be identified, namely the basal side of the epithelium and the apical junctional network (**Figure 6A-B**). Both surfaces are mesh-like structures of high inclination, high signal intensity and high contrast. The vesicle lumen also contains cell debris of high intensity and contrast that are not part of any surface of interest.

**FIGURE 6.**
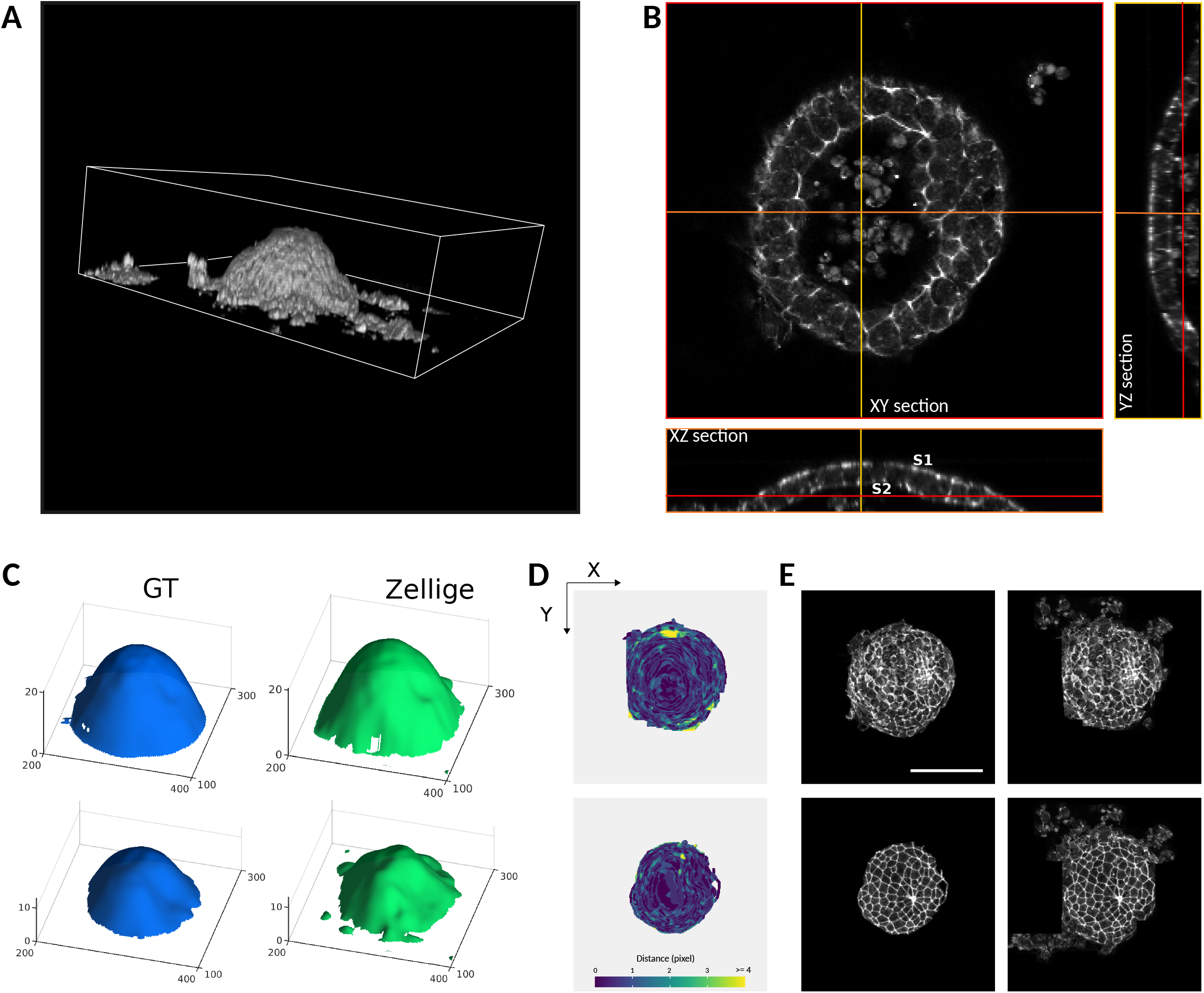
(A-B) Volume rendering (A) and orthogonal sections (B) of a confocal 3D image of a (half of) inner ear organoid, which has been fixed and stained with phalloidin to reveal F-actin. The dataset (of dimensions 520 × 465 × 35 pixels) includes two dome-shaped epithelial surfaces of interest, forming the apical (inward) and basal (outward) sides of the organoid. (C) 3D representations of the height map extracted by Zellige (in green) and the GT height map (in blue), of the epithelium surface. (D) Color-coded error maps of the reconstructed height-maps for the apical (left) and basal (right) epithelial surfaces of the organoid. The surfaces of interest are reconstructed with an error of < 2 pixels over a large majority (96% and 93% for the apical and basal surfaces, respectively) of pixels, as well as on average (RMSE ~ 0.8 and 1.1 for the apical and the basal surfaces, respectively). (E) Projections localized to the GT height-maps of the epithelium surface (panels on the left), and the height-maps extracted by Zellige (panels on the right). Scale bar 100 μm.

**Figure 6C** shows a 3D graphical representation of the height-maps reconstructed by Zellige and those reconstructed by an expert biologist, taken as ground truth (GT). Due to their dome-shaped topography, the manual segmentation of these surfaces was rather laborious, and is more likely prone to errors in the regions of high inclination. Despite this, we observe an excellent match between the two reconstructed and corresponding GT height-maps (**Figure 6D**). The distance between the two height maps is <2 for the large majority (> 92%) of pixels, while larger error values occur locally in regions of near-vertical inclination of the surfaces. The RMSEs of both apical and basal surfaces are close to 1, showing that on average the reconstructed height-maps match the corresponding GT with about pixel accuracy. The coverage of the reconstructed height-map is nearly 100% for the basal surface, and ~88% for the apical surface. **Figure 6E** shows the 2D projections of the 3D image obtained for each of the reconstructed surfaces, as compared to the projected GT height maps. Thus, for the extraction for these highly inclined surfaces, Zellige produces height maps of a quality comparable to those obtained from manual expert segmentation.

In this example, Zellige could be combined with a 2D cell tracking framework such as TissueMiner [3] to perform cell dynamics analysis. Note that geometric distortions introduced in the projected surfaces by the epithelium inclination could be corrected for, using complementary tools such as DProj [11]. This approach could provide a means to quantitatively address how an inner ear organoid epithelium patterns at the cellular and organoid scales, while quantifying the epithelial thickness changes due to cellular intercalation or cell shape changes in the depth of the epithelium. This would also permit to better characterize the variability of inner ear organoids within in a given aggregate, and it could allow one to explore how the organoid interacts with surrounding tissues and how these interactions influence the differentiation of their constituent sensory cells.

#### A sensitivity analysis reveals the robustness of Zellige in extracting surfaces from biological images

To evaluate further the quality and robustness of the segmentation obtained by Zellige, we carried out a sensitivity analysis of the reconstruction on each of the samples tested. This analysis consisted in varying one control parameter at a time (**Figures S2-S6**, **Figure 7**, Supplemental note 4), while keeping the other parameters fixed at a nominal value (Supplementary Table S1). The RMSE and coverage of each of the reconstructed surfaces were evaluated and plotted as a function of the value of the parameter that was varied.

**FIGURE 7.**
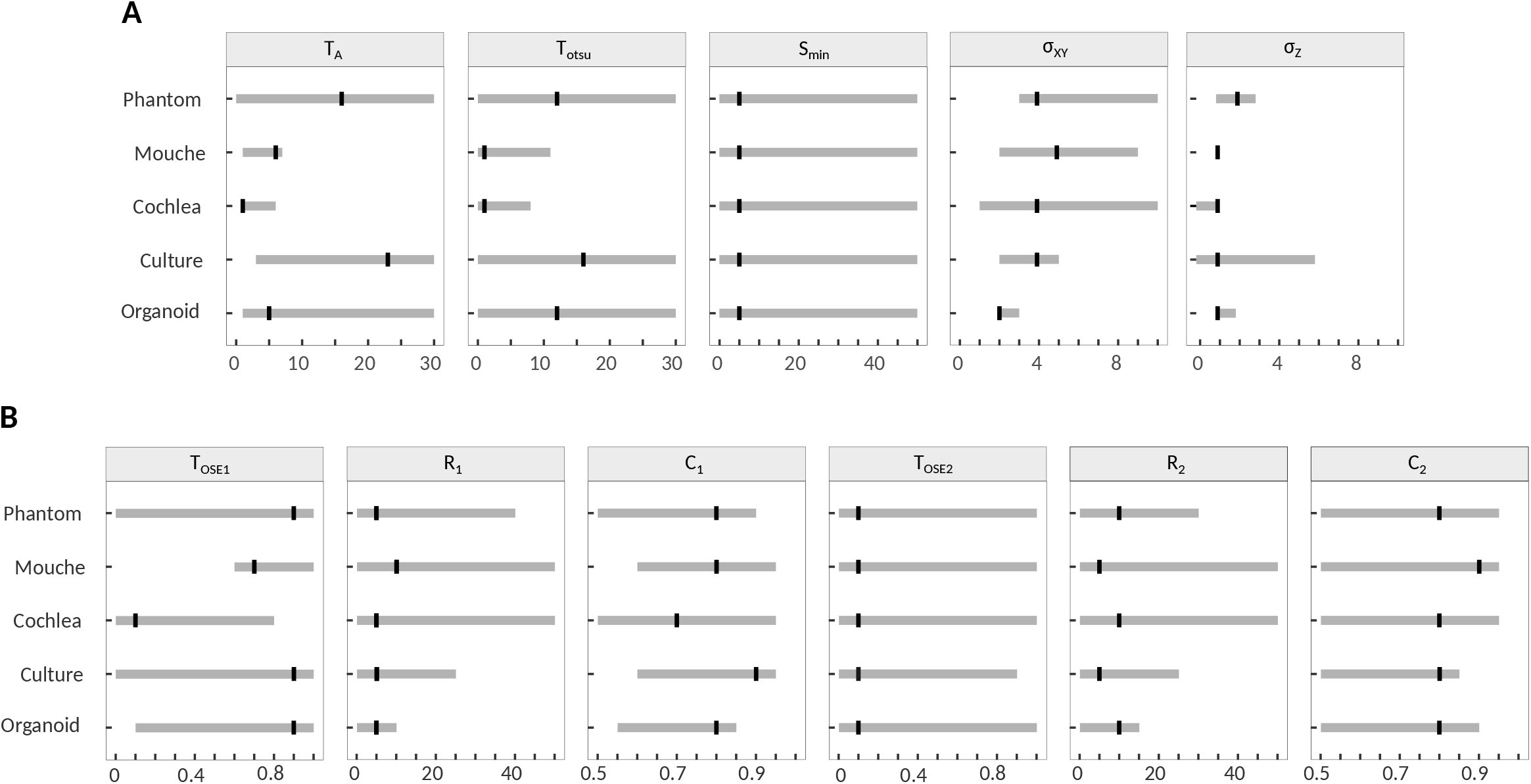
Summary of the sensitivity analysis. The intervals indicated in grey for each parameter and each of the images tested correspond to the parameter values for which the reconstruction satisfies high quality criteria defined by RMSE ≤ 1.5 and coverage ≥ 85%. Black marks indicate the reference value obtained by manual adjustment for each image (cf. Supplemental note 2). (A) Parameters of the surface selection step. (B) Parameters of the surface assembly step.

**Figure S3** shows the results of the sensitivity analysis carried out on the image of **Figure 3** (Example 1, pupal fly specimen), when varying the parameters controlling step 1, *i.e.* the selection of putative surface pixels (parameters *T*_A_, *T*_otsu_, *S*_min_, *σ*_xy_ and *σ*_z_). As can be seen, the variations of the two classification threshold parameters *T*_A_, *T*_otsu_ and of the minimum island size *S*_min_ in their respective intervals does not substantially modify the RMSE and the coverage of the reconstructed surfaces, whose values remain roughly constant for *T*_A_ ≤ 8, *T*_otsu_ ≤ 12 and over the entire *S*_min_ interval (**Figure S3A**). Within these intervals, the RMSEs of all the reconstructed surfaces remain ≤1.5, while the coverage values are > 95% for surfaces S1 and S2, and > 85% for surfaces S3 and S4. The surfaces S1 and S3 show lower signal intensity and lower contrast than surfaces S2 and S3, making them more difficult to extract. Surface S4 has the lowest contrast, and fails to be reconstructed if the classification threshold values are too stringent (namely for *T*_otsu_ > 12, or *T*_A_ > 8). Nevertheless, the intervals of stability of *T*_A_ and *T*_otsu_ (that is, the intervals of values over which a high-quality extraction of all surfaces is obtained) remain relatively wide (cf. **Figure 7**). The smoothing parameters *σ*_xy_ and *σ*_z_ also have some effect on the reconstruction of the surfaces. When *σ*_xy_ is less than 3, the RMSE is higher for the mesh-like epithelial surfaces S2 and S4 (formed by the junctional network of the epithelium). A minimal smoothing along the axial direction is also important to ensure that the reconstructed surfaces are not too fragmented, preventing their complete reconstruction. Yet, *σ*_z_ should not be chosen too large either, to avoid merging nearby surfaces along the z axis. In this case, the closely positioned surfaces S1 and S2 are well-separated if setting *σ*_z_ close to 1, but they become merged when *σ*_z_ > 2. In general, the surface construction parameters have little effect on RMSE and coverage (**Figure S3B**). The sensitivity analysis on this challenging specimen shows a good robustness of Zellige to extract the four surfaces with a single set of parameters, each of which can be chosen in a reasonably wide interval considering the other fixed.

The results of the sensitivity analyses performed with the other biological image stacks (examples 2, 3 and 4 described above) are shown in **Figures S4**, **S5** and **S6** (Supplemental note 4). These results are summarized in **Figure 7**, which show the stability intervals over which the extracted height-maps satisfy the criteria RMSE ≤ 1.5 and coverage ≥ 85%, for which the reconstruction can be considered of high quality. Overall, the stability intervals for the two classification threshold parameters (*T*_A_ and *T*_otsu_) are narrower for specimens containing a surface of low signal intensity and low contrast, while still covering about 1/4 of their respective width. The graphical user interface of Zellige allows the user to adjust *T*_A_ and *T*_otsu_ interactively, making it intuitive to search for reasonable values. We found that 2 ≤ *σ*_xy_ ≤ 3 and *σ*_z_ = 1 generally give high quality results for all tested specimens (**Figure 7A**). We therefore expect only little adjustment to be required by the user on the smoothing parameters from their default permissive values (set to *σ*_xy_ = 1 and *σ*_z_ = 1). Values of *σ*_z_ that are too large may lead to the merging of a surfaces with a nearby structure of high contrast (surface or else), as it happens for the epithelium surface in the cochlea specimen, which merges with the underlying mesenchyme signal when *σ*_z_ is greater than 2 **(Figure 4 and Figure S4).** The effect is even more pronounced for surfaces of high inclination (**Figures 2, 6** and **Figures S2, S6**), or presenting a particularly rough texture (**Figures 5** and **Figures S5**).

Regarding the control parameters of step 2 (the assembly step), for the four examples presented except for example 1, the quality of the reconstructed surfaces is stable and high over the main part of their respective intervals of variation, deteriorating occasionally only when extreme values (≃ 1) are used for these parameters (see **Figures S2-S6, Figure 7B** and Supplemental note 4). Example 1 poses particularly stringent constraints on the control parameters of the reconstruction due to the requirement of reconstructing the 4 surfaces of different contrast and texture present in this sample using the same set of parameters.

Finally, the computation times for running Zellige on a given dataset ranged between a few seconds and a minute on a standard PC computer (see **Figure S7** and Supplementary note 5), except in a few exceptional cases corresponding to extreme values of the control parameters. As a safeguard, a stopping criterion could be implemented so as to exit the run (declaring the current parameter values invalid) if the surface assembly computation exceeds a user-prescribed duration.

Overall, the sensitivity analysis indicates that the surface extraction performed by Zellige is robust to variations of the control parameters of step 1 (surface pixel selection step). In general, the reconstruction is more sensitive to the amplitude threshold parameter (*T*_A_), and the Otsu threshold parameter (*T*_otsu_) should be kept sufficiently low for samples containing surfaces of intensity close to the background. However, in some cases such as in our example 3, the opposite is true. Thus, the two threshold parameters play somewhat complementary roles, and the possibility to adjust them independently is useful in practice to be able to cover as many cases as possible. A smoothing along *xy* appears necessary to correctly reconstruct the surfaces supported by a junctional mesh. Not surprisingly, best results are obtained when the radius of the gaussian filter used for this (parameter *σ*_xy_) is adapted to the mesh (or cell) size. Likewise, a smoothing along z is beneficial, but the extent of this smoothing (parameter *σ*_z_) should not be too large to avoid causing the fusion of nearby surfaces. With a few exceptions, the values of the RMSE and coverage show little sensitivity to the values of the parameters controlling step 2 (surface assembly step), at least once putative surface pixels have been properly selected. In the presence of several surfaces of potentially very different sizes, the parameter controlling the fraction of OSE sizes allowed for OSE seeds (parameter *T*_OSE1_) should be relatively large (≥ 0.5 or greater, *i.e.* allowing more than 50% of the largest OSE sizes for seeds) to allow the extraction of a surface of small size (for example, to extract the surface n° 4 of example 1, which covers less than 20% of the xy-field of view, *T*_OSE1_ must be larger than 0.6). Extreme values (close to 1) for the connectivity rates (C1 and C2) are too stringent and lead to a drop in the coverage of the reconstructed surfaces. To sum up, we see that the most critical parameters for a satisfactory extraction of the different surfaces are those controlling step 1. In most cases the parameters controlling step 2 do not need to be adjusted and can be fixed to their default reference values.

## Conclusion

We have developed Zellige, a new tool to extract multiple surfaces from 3D fluorescence microscopy images. Zellige automatically finds surfaces by first identifying putative pixels that are likely to belong to a biological surface, and second by assembling a surface through connection of adjacent pixels satisfying natural proximity constraints. By using Zellige on synthetic epithelium images we have shown that it accurately reconstructs a surface with excellent performances in terms of both the distance to the ground-truth height-map and the surface coverage (**Figure 2**). Zellige can deal with complex images containing multiple surfaces, with computation times not exceeding a few tens of seconds on a standard computer. Importantly, the user is not required to specify the number of surfaces to be extracted. In the *Drosophila* specimen (**Figure 3**), the software readily extracts the 4 surfaces of interest that could be identified. Since Zellige detects putative surface pixels in the first step by combining local and global thresholds, it can deal with images where the multiple surfaces display different features, such as in the mouse cochlear embryo (**Figure 3**). With this difficult dataset, we could also confirm Zellige’s robustness against very low signal-to-noise levels. The constructive approach of surface region growing used by Zellige in its second step enables it to circumvent the surface smoothness requirement, that is classically assumed by other surface extraction tools. For instance, it could reconstruct the highly irregular surface of a bronchial tissue infected by SARS-CoV-2 (**Figure 5**).

The robustness and flexibility of Zellige come at a price, namely, the requirement to specify 12 parameters when running the surface extraction. However, the sensitivity analysis we performed shows that adjusting only 4 of these parameters is enough in practice to handle a wide range of image types. These parameters correspond to intuitive notions *(e.g.,* thresholding and smoothing), which makes Zellige particularly easy to use. The Fiji interface that we implemented to perform this adjustment should make Zellige even more user-friendly and effective for biological applications.

To our knowledge, Zellige is the only open-source tool that can extract an unspecified number of epithelial surfaces from a 3D volume, possibly larger than two. This unique feature is especially useful in complex images that could be processed only by specialized tools before. For instance, Zellige can extract surfaces with projections on the *xy* plane that completely overlap, such as the basal and apical epithelia in the organoid image of **Figure 6**. Previously, such an image could be processed only by tools that relied on segmenting a mesh around the object surface, such as MorphoGraphX or ImSAnE [16, 47].

The flexibility and robustness of Zellige should allow to considerably relax the constraints that were previously imposed on the sample preparation and the image acquisition steps by the subsequent analysis. Indeed, Zellige can accommodate any number of surfaces in the acquired volume, overlapping or not, and of different contrast features. Zellige also showed excellent robustness against image noise. This should make it particularly useful in imaging contexts that are not easily amenable to automated analysis, such as intravital imaging. Finally, it is worth noting that Zellige is a generalist and modular method. With some adaptation of the surface pixel selection step, it could be used with imaging modalities beyond the scope of this article, for instance, in extracting the irregular and noisy surfaces of biological objects imaged with 3D electron microscopy images.

## Methods

### Implementation

Zellige was devised with the goal of achieving accurate segmentation of multiple biological surfaces from 3D confocal images. Unlike other existing surface extraction tools, it makes no assumption on the number of surfaces to be extracted and does not require the surfaces of interest to be the structures of highest contrast in the image. Zellige is written in Java, relying on the ImgLib2 library [48] and is distributed as a Fiji plugin, with a graphical user interface (GUI) designed to allow users to quickly find a good set of extraction parameters for a given image.

Zellige extracts each surface present in the image in the form of a height-map (or *z*-map), that is, a mapping

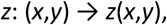

which associates to each point (*x,y*) over which the surface projects, the z-coordinate of the unique pixel (*x,y,z*) belonging to the surface. Each extracted height-map is then used to produce a 2D projection of the 3D stack restricted to a small sub-volume (of user-selected width) centered around the corresponding surface. To achieve a robust extraction, Zellige proceeds in two algorithmic steps, which are only outlined below (see Supplementary note 1 for implementation details).

In the first step, or *surface pixel selection* step, a segmentation is applied to the 3D image to select pixels that likely belong to a surface of interest (**Figure 1 and Figure S1A**). These putative surface pixels are detected as local maxima of image intensity along the z-axis, after using two independent binary classifiers, one based on pixel contrast and the other one on pixel intensity. Five adjustable parameters control the selection step: two threshold parameters (*T*_A_ and *T*_otsu_) control the strength of the binary classifiers applied on contrast and intensity, respectively, and three parameters (*S*_min_, *σ*_xy_ and *σ*_z_) control clean-up operations applied at the end of the classification (removal of small isolated spots, and local averaging along the *xy* plane and the z axis, respectively).

In the second step, or *surface assembly step,* an iterative algorithm is used to extract the heightmaps of each of the surfaces present in the image (**Figure 1 and Figure S1A**). The assembly starts by grouping neighboring putative surface pixels together within each orthogonal (*xz* or *yz*) section of the 3D image, in order to form a set of building blocks referred to as *orthogonal surface elements* (OSEs). These building blocks are then used to assemble the surfaces, in a process analogous to jigsaw puzzles, where OSEs adjacent to the surface boundary are added if they match this boundary, and rejected otherwise, until no matching OSE can be found (**Figure S1**). In order to increase the robustness of the assembly step, Zellige applies it in two rounds, proceeding along different axes during the first and second rounds. Each round is controlled by 3 adjustable parameters: a threshold parameter (0 ≤ *T*_OSE_ ≤1) sets a minimum size for the building blocks that can be used as seeds to initiate the assembly of a surface; and two other parameters (0 ≤ *R* ≤ 50 and 0 ≤ *C* ≤ 1) set the matching constraints used to accept or reject the addition of OSEs to a surface. The assembly step is thus controlled overall by 6 parameters, *i.e.* two groups of 3 parameters (*T*_OSE1_, *R*_1_, *C*_1_) and (*T*_OSE2_, *R*_2_, *C*_2_) controlling the first and second assembly rounds, respectively.

Finally, the height-maps of each of the reconstructed surfaces are used to obtain a corresponding 2D projection (**Figure 1**). In practice a maximum projection restricted to a subvolume of width δz (δz being a user-defined parameter) centered around the surface of interest is performed.

### Availability of data and code

The project homepage below contains the source code, installation instructions, and documentations.

### Zellige

- Project name: Zellige.
- Project homepage: https://gitlab.pasteur.fr/ida-public/zellige-core
- URL for the Fiji plugin (Update > Manage update sites): https://sites.imagej.net/Zellige/
- Gitlab Branch: master
- Operating systems: Platform independent.
- Programming language: Java.
- Compiled in Java8
- Other requirements: Runs from Fiji [28].
- License: BSD 2
- Any restrictions to use by non-academics: None.

### Scripts to create Phantoms

- Project name: Phantoms.
- Project homepage: https://doi.org/10.5281/zenodo.6414596
- Operating systems: Platform independent.
- Programming language: MATLAB.
- License: BSD 2
- Any restrictions to use by non-academics: None.

### Data sets

Image 3D stacks, ground-truth height map and height maps produced by Zellige are available on Zenodo under the CC-BY license: https://zenodo.org/communities/zellige/ [29, 33, 42, 44, 46].

### *Human* bronchial epithelium imaging

The data used in Figure 5 were taken from the recent study [43] to which we refer for the preparation and imaging of human bronchial epithelium cultures. In brief, MucilAirTM were purchased from Epithelix (Saint-Julien-en-Genevois, France) and cultured for at least 4 weeks to reconstruct a differentiated human bronchial epithelium *in vitro* and stained as previously described. Images of the cultures were acquired using an inverted Zeiss LSM 710 confocal microscope controlled by the ZEN pro 2.3 software. Z-stack images of whole-mount samples were acquired with a Zeiss Plan Apochromat 63x oil immersion lens (NA=1.4). The image used here was published in Robinot *et al.* [43] under the CC-BY-4 license.

### *Drosophila* imaging

Flies were raised at 25°C under standard conditions. Pupae were collected for imaging as described previously [49]. Ecad::GFP flies [50] were used for live imaging as previously described [1]. In brief, images were acquired with a spinning disk microscope from Gataca Systems driven by the MetaMorph software. The system is equipped with an inverted Nikon TI2E stand, a motorized XYZ stage, and a Nikon Plan Apo 60x oil immersion (NA=1.4) lens and with a Prime95B camera.

### Cochlea imaging

The inner ears from wild-type (C57BL/6) mice were rapidly dissected from temporal bones at embryonic stages E14.5 in HEPES-buffered (10 mM, pH 7.4) Hanks’ balanced salts solution and fixed in 4% paraformaldehyde, 1 hour at room temperature. Specimen were permeabilized and stained for phalloidin-Atto 565 (Sigma) as previously described [39]. Fluorescence images were obtained with a swept-field confocal microscope (Opterra II) from Brucker. This system is equipped with a Nikon Plan Fluor 60x oil immersion lens (NA=1.4).

Biologists in this study hold a designer certificate of animal experimentation (level 1), allowing them to perform experimental work on animals in strict accordance with the European directive 2010/60/EU, and French regulations. The Ethics Committee of the Institut Pasteur (Comité d’Ethique en Experimentation Animale – CETEA) has approved this study with the project identifier dha170006. This approval is based on careful compliance to the 3Rs principle in the care and use of animals (Annex IV – 2010/60/EU).

### Inner ear organoid imaging

ESCs derived from blastocyst-stage embryos of R1 mice (mESCs) (ATCC, SCRC-1036) were maintained in feeder-free culture on 0.1% w/v gelatin (Sigma) coated substrates using LIF-2i medium as established previously [51]. The organoids were generated following the previously published protocol [45, 51]. Aggregates were harvested at day 14 and fixed in 4% v/v PFA (Electron Microscopy Sciences) overnight at 4°C. After blocking (PBS; 10% v/v normal goat serum; 0.1% v/v Triton X-100), the aggregates were stained for phalloidin Atto 565 (1:1000) (Sigma) overnight at RT on a shaker, and washed three times with PBS containing 0.1% v/v Triton X-100 for 1 h each at RT. Prior to imaging, the aggregates were incubated in a modified version of ScaleS solution containing 4 M Urea (Sigma), 40% w/v D-Sorbitol (Sigma), and 0.1% v/v Triton X-100, for 3-5 days to clarify the tissue. Finally, the aggregates were whole-mounted using the ScaleS solution and imaged using a confocal laser scanning microscope (A1R HD25, Nikon) equipped with a Nikon 25x silicon oil immersion lens (NA=1.05).

## Acknowledgements

We thank Maia Brünstein of the Hearing Institute Bioimaging Core Facility – C2RT/C2RA and Florian Rückerl of the Photonic BioImaging – C2RT/C2RA, for sharing their expertise on light microscopy. We thank Romain Levayer for sharing flies and lab space with us. We thank Gilles Trébeau (https://fr.linkedin.com/in/gilles-trebeau-427601212) for helpful discussions regarding the Java frontend deployment of Zellige. We thank Lisa Chakrabarti, Rémy Robinot and Vincent Michel for sharing data. This work was supported by *Fondation pour l’Audition* (FPA-IDA-STARTING-GRANT); *Institut Pasteur* (PTR#272); DIM ELICIT’s grant from *Région Ilede-France* (QP-IDF-DIM-ELICIT-2019); The French *Agence Nationale de la Recherche* (ANR-21-CE13-0038-013-01 and LabEx LIFESENSES ANR-10-LABX-65).

