## Supplemental Note for "Extracting multiple surfaces from 3D microscopy images in complex biological tissues with the Zellige software tool"

### Supplemental note 1: Overview of Zellige implementation

Zellige takes as its input a 3D confocal fluorescence microscopy image $I(x,y,z)$ containing one or several surfaces of interest. The aim of the software is to segment all surfaces present in the image, and to extract them in the form of height-maps (**Figure 1**). To this end, Zellige proceeds through two main computational steps. In step 1, the software performs a selection of the putative surface pixels, *i.e.* pixels that likely belong to a surface within the image (**Figure S1A, upper panel**). These putative surface pixels are detected as local maxima of image intensity along the z-axis, after using two independent binary classifiers, one based on pixel contrast and the other one on pixel intensity ('amplitude' and 'Otsu' classifiers, described below). The output of step 1 is a binary classification masks $B(x,y,z)$, which defines the set of selected putative surface pixels in the image volume. In step 2, Zellige performs the reconstruction of the surfaces, taking as input the array $B(x,y,z)$ and using an assembly algorithm analogous to a jigsaw puzzle construction (Figure S1A, lower panel). The main output of step 2 is a list of the height-maps $h_1,\ldots,h_n$ of each of the surfaces that have been assembled. Finally, the projections of the image $I$(x,y,z) localized near each of these height-maps are obtained. The two computational steps of Zellige (step 1 and step 2) are described below in more details.

***Step 1 – Selection of putative surface pixels (Figure S1A, upper panel).***

After filtering the image by a 3D gaussian blur (with standard deviations of 2x2x1 pixels) to reduce noise artifacts, putative surface pixels are detected as the local maxima of image intensity along the z-axis, that is, the pixels ($x,y,z$) where the partial derivative $\partial I/\partial z$ of image intensity with respect to $z$ changes sign from positive to negative along the z-axis. However, to decrease the number of false positive among these local maxima, the image is first subjected to a classification scheme combining two independent binary classifiers: the 'amplitude' and the 'Otsu' classifiers.

The amplitude classifier is based on a local contrast indicator that quantifies the saliency of a local intensity maximum detected along the z-axis. Specifically, an amplitude $A(p)>0$ is assigned to each such maximum $p=(x,y,z)$, defined as the peak-to-peak difference between the intensity of the given local maximum $p$ and the intensities of the nearest local intentity minima $p_1$, $p_2$ along the z-axis, namely, $A(p) = \max(I(p)-I(p_1),I(p)-I(p_2))$ (see **Figure S1B**). An zero amplitude is assigned to all the other (non-local maxima) pixels, and the resulting array $A(x,y,z)$ is normalized to have values between 0 and 255. A binary classifier mask $B_A(x,y,z)$ is then defined by setting $B_A(x,y,z) = 1$ if $A(x,y,z) \geq T_A$, and $B_A(x,y,z) = 0$ otherwise, where $0 \leq T_A \leq 255$ is an adjustable threshold setting the minimum allowed salience of a local maximum to be selected as a putative surface pixel.

The Otsu classifier is based on the Otsu method, a popular method of segmentation of an image into two classes of "background" and "foreground" pixels [1]. In Zellige, a set of local Otsu thresholds are estimated on a grid of subvolumes of size 40x40x3 subdividing the image stack. These thresholds are regrouped and smoothed using a 3D gaussian filter (with standard deviations of 5x5x1 pixels) to produce a 3D-array of local Otsu thresholds $T_{otsu,loc}(x,y,z)$. An adjustable global Otsu threshold $T_{otsu}$ is also applied to remove low-lying background pixels. The Otsu binary classifier mask $B_{otsu}(x,y,z)$ is then defined by setting $B_{otsu}(x,y,z)=1$ if $I(x,y,z) \geq \max(T_{otsu,loc}(x,y,z),T_{otsu})$, and $B_{otsu}(x,y,z)=0$ otherwise.

The two classifiers just described are combined by taking the product (or intersection) mask $B(x,y,z) = B_A(x,y,z)B_{\text{otsu}}(x,y,z)$, which defines a pre-selected set of putative surface pixels. This mask is further processed by additional clean-up operations (**Figure S1A upper panel)** designed to i) remove small isolated clusters or 'islands' formed by putative surface pixels unlikely to belong to a surface of interest, and ii) to partially fill possible surface holes by applying a final gaussian blur to the mask. In the first operation (island removal), an island is defined as a 4-connected component formed by putative surface pixels within some *xy*-section of the image, which is of a size smaller or equal than a threshold $S_{\min}$, and is not connected to any other selected maxima lying on one of the neighboring z+1 or z-1 sections. The minimum tolerated island size $S_{\min}$ is an adjustable parameter set by default to 5 pixels. Any cluster of a size $\leq S_{\min}$ is removed from the list of surface pixels. The second operation (gaussian blur) depends on two adjustable parameters, namely the gaussian blur radiuses $\sigma_{xy}$ and $\sigma_z$ used along *xy* and *z*, respectively. After the smoothing, a detection of local maxima along z is performed once more on the mask $B(x,y,z)$, this time without classification; this results in a final binary detection mask $M(x,y,z)$ containing the set of putative surface pixels to be used in step 2.

***Step 2 - Surface reconstruction by assembly of orthogonal surface elements (Figure S1A, lower panel).***

The assembly of surfaces implemented in Zellige proceeds in two phases. First, within each orthogonal (*xz* or *yz*) section of the 3D image, building blocks called *orthogonal surface elements* (OSEs) are constructed out of the selected surface pixels. The OSEs correspond to contiguous chains of pixels delineating the profile of some surface intersecting the given orthogonal section. Formally, they are defined as the connected components of a graph $G_{OSE}$ defined on the set of putative surface pixels contained within each orthogonal section, with a neighbor structure imposing contiguity constraints assumed to hold for such a profile (**Figure S1C,D**).

The OSEs then serve as base elements in the subsequent process of surface assembly. This process starts with a *surface seed*, defined as an OSE whose size is ranked before the $T_{OSE}$'th percentile (where $0 \leq T_{OSE} \leq 1$ is some adjustable fraction) among the list of OSE sizes ranked in decreasing order (in other words, only the OSEs of sizes ranked among the largest $T_{OSE}$ percents can be used for a surface seed). Starting from a seed, a surface $S$ is assembled sequentially by adding new adjacent OSEs provided they fulfill some graph-matching condition. Briefly, a candidate OSE $\sigma$ adjacent to the boundary of $S$ is tested by computing two quantities, the overlap $R(S,\sigma)$ and connectivity $C(S,\sigma)$ of $\sigma$ relative to $S$, which measure how well the shape of $\sigma$ matches to the surface boundary (**Figure S1E**). The OSE $\sigma$ is then added to the surface if its overlap and connectivity are greater than some adjustable thresholds $R_0$ and $C_0$. The surface $S$ is complete when no OSE matching its boundary can be found. A new surface is then assembled, by choosing a new OSE seed that did not belong to $S$, and reiterating the above construction. The whole assembly process ends when there is no OSE seed left which does not belong to some surface already constructed. At this point, a list of $n'$ reconstructed surfaces $S_1,\ldots,S_{n'}$ has been obtained, not all of which will be retained, however: Surfaces that are too small to be genuine surfaces of interest are first removed; the minimum surface size is set by another adjustable parameter $0 \leq p_{min} \leq 1$, which defines the minimum fraction of the image's *xy* field of view that should be covered by a reconstructed surface. In addition, it may happen that two or more among the remaining surfaces are identical, except for a small fraction of the OSEs of which they are composed. When for a pair of surfaces this fraction is smaller than some tolerance value, the surfaces are considered redundant and merged, resulting in one multi-valued height-map in which some positions $(x,y)$ correspond to more than one $z$-value. After this pruning and merging phase, one is left with a list of $n<n'$ distinct but possibly multi-valued surfaces $S_1,\ldots,S_n$ .

To increase the robustness to the above construction, Zellige applies it in two rounds: in the first round, OSEs are constructed within *xz* orthogonal sections, and surfaces are assembled along the *y* axis using relatively loose assembly parameters ($T_{OSE1}$, $R_1$, $C_1$). In the second round, each of the surfaces $S_1,\ldots,S_n$ assembled in the first round is reconstructed in turn, keeping only the pixels of a given surface. This time however, the OSEs are constructed within yz sections, and the surfaces are assembled along the *x* axis using more stringent matching constraints ($T_{OSE2}$, $R_2$, $C_2$, with $T_{OSE2} < T_{OSE1}$, $R_2 > R_1$, and $C_2 > C_1$).

At the end of the two assembly rounds, the ambiguities still present in the surfaces $S_1,\ldots,S_n$ are removed to form a final list of genuine (mono-valued) height-maps $h_1,\ldots,h_n$. This is done for each surface $S_i$, by replacing the height-values of each multiple point, *i.e.* each position $(x,y)$ at which two or more surface points have been detected, by a single value $h_i(x,y)$, or by no value at all, as follows: i) If two points have been detected at heights $z_1$ and $z_2$ that differ by less than 2 (i.e. such that $|z_1-z_2| \leq 2$) in pixel units, these two values are averaged by setting $h_i(x,y)=(z_1+z_2)/2$; iii) In all the other cases, i.e. if more than 2 points have been detected, or if two points have been detected whose heights $z_1$ and $z_2$ differ by more than 2, the points detected in Si at $(x,y)$ are discarded, and no value is ascribed to the height-map $h_i$ at that point.

### Supplemental note 2: Generation of phantom 3D images

The synthetic images used in this project were generated in MATLAB (The MathWorks) using custom code available our public Gitlab instance: . The phantom 3D image shown in **Figure 2** models a typical stack of confocal images of epithelial and non-epithelial structures. It contains three superimposed surfaces, each constructed using a simple algebraic formula. The surfaces generated are of two types: full surfaces, presenting a homogeneous signal over the entire surface, or surfaces presenting a signal restricted to a polygonal mesh mimicking the mesh of apical cellular junctions of an epithelium observed at its surface. To generate these models, we constructed a height-map $z = h(x,y)$ and a projected image $i_0(x,y)$, both having the same format $W \times H$ (with $W = H = 512$ or 1024). For a full surface, the projection is taken constant, $i_0(x,y) = 1$. For a surface restricted to a mesh, the image $i_0(x,y)$ is defined as the mask of a Voronoi mesh constructed on a set of $N$ points ($100 \leq N \leq 300$) distributed at random in the field of the image. The maps $h(x,y)$ and $i_0(x,y)$ are then used as ground truth (GT) to generate a non-noisy 3D image $I_0(x,y,z)$ with ($0 \leq x \leq W - 1$, $0 \leq y \leq H - 1$, $0 \leq z \leq D - 1$) in which a discretized version of the surface is inserted, according to the definition: $I_0(x,y,z) = i_0(x,y)$ if $z = [h(x,y)]$, $I_0(x,y,z) = 0$ otherwise. The image $I_0(x,y,z)$ is then subjected to 3D filtering by a Gaussian filter of a size reflecting the point spread function under the sampling conditions typical of confocal images (standard deviations $2 \times 2 \times 3$ pixels along $x$, $y$ and $z$). Finally, a Poissonian random noise is added to the result (modeling the photon noise of the confocal images) to produce a final image $I(x,y,z)$ having a given signal-to-noise ratio (SNR).

### Supplemental note 3: Comparing the reconstructed and ground truth height-maps

#### Definition of the ground-truth (GT) height-map

To evaluate the ability of Zellige to reconstruct surfaces present in a 3D image, we define a ground-truth (GT) height-map, that associates each pixel $(x,y)$ of the surface to its true position in the axial (z) dimension. In the case of phantom images, this GT height-map is known exactly. In the case of 3D biological microscopy images, they were obtained as described in [2] by manual annotation, using the Icy software [3]. In brief, the GT height-maps were generated by first manually marking the regions containing the surface of interest on each $z$ section of the image stack. The localization of the surface in z can be ambiguous, which may lead to the selection of multiple $z$ values for a single $(x,y)$ position. When this is the case, the different $z$ values are averaged.

When more than one surface of interest was present in the original 3D image, we repeated the above manual segmentation procedure so as to generate one GT height-map for each surface. The height-map of a given surface of interest reconstructed by Zellige or some other surface extraction software could then be compared quantitatively to the corresponding GT height-map, by computing its root mean square error and its coverage.

##### Root mean square error of the reconstructed height-map

The Root Mean Square Error (RMSE) is a standard measure of the error made by an estimator of a continuously varying quantity. This error is defined here as the root mean square (or Euclidean) distance between a reconstructed height-map $h(x,y)$ and the ground truth height-map $h_{GT}(x,y)$. Note that neither of these height-maps necessarily covers the entire $xy$ field of the image. In general, one or both of the values $h(x,y)$ and $h_{GT}(x,y)$ remain undefined for some points $(x,y)$, because no corresponding surface point has been detected or selected. We denote by $n_T$ the number of pixels $(x,y)$ for which both the height-maps $h(x,y)$ and $h_{GT}(x,y)$ are defined. In the Zellige implementation, a height-map takes >0 values on the pixels where it is defined, and the value 0 on the pixels where it is undefined. With this convention, the RMSE is defined by the formula:

$$\mathrm{RMSE} = \{1/n_T \sum_{(x,y)} |h(x,y) - h_{GT}(x,y)|^2\}^{1/2},$$

where the sum runs over the set of pixels for which $h(x,y)$ and $h_{GT}(x,y)$ are both > 0. A good quality standard for an efficient surface reconstruction algorithm is to produce a height-map with an RMSE of less than 1 (in pixel units), corresponding to an average accuracy of a fraction of a pixel, i.e. typically subpixel distances between the reconstructed and the GT height-maps.

##### Coverage of the reconstructed height-map

The coverage of the reconstructed height-map (relative to the corresponding GT height-map) is another measure that must be considered when judging the quality of surface reconstruction. It is defined here as the ratio

$$\mathrm{Rec} = n_T/n_{GT}$$

where $n_{GT}$ is the number of pixels of the GT height-map.

#### Supplemental note 4: Sensitivity Analysis

##### Sensitivity analysis by means of a one-parameter scan

A sensitivity analysis provides an important test of a surface reconstruction algorithm, by quantifying the influence of the various control parameters on the quality of the reconstruction. The aim here is to determine if and when a small change in the value of a given parameter might have a big impact on the RMSE and the coverage of the reconstructed surfaces. This analysis proceeds by scanning each control parameter in turn (one at a time) over its entire working range of values. For each value tested, the surface reconstruction algorithm is run, and the RMSE and coverage of the reconstructed surface(s) of interest are computed. As a result of this analysis, the parameters which have the most critical influence on the results can be identified, as well as those which have little or no effect. The sensitivity analysis carried out in this study was performed for each control parameter of Zellige in succession: parameters ($T_A$, $T_{otsu}$, $S_{min}$, $\sigma_{xy}$, and $\sigma_z$) of the surface pixel selection step, and parameters ($T_{OSE1}$, $R_1$, $C_1$, $T_{OSE2}$, $R_2$, $C_2$) of the surface assembly step. Results obtained on the phantom image of **Figure 2** and on the biological 3D images of **Figures 3-6** are described below (**Figures S2-S6**), and summarized in **Figure 7**.

| | | Selection parameters | | | | | Construction parameters | | | | | | | |
|---|---|---|---|---|---|---|---|---|---|---|---|---|---|---|
| | **Surface** | **TA** | **Totsu** | **IS min** | **σXY** | **σZ** | **TOSE1** | **R1** | **C1** | **TOSE2** | **R2** | **C2** | **RMSE** | **Coverage** |
| **Phantom** | n°1 | 16 | 12 | 5 | 4 | 2 | 0.9 | 5 | 0.8 | 0.1 | 10 | 0.8 | 0.23 | 1.00 |
| | n°2 | | | | | | | | | | | | 0.31 | 1.00 |
| | n°3 | | | | | | | | | | | | 0.56 | 1.00 |
| **Fly** | n°1 | 6 | 1 | 5 | 5 | 1 | 0.7 | 10 | 0.8 | 0.1 | 5 | 0.9 | 0.93 | 0.99 |
| | n°2 | | | | | | | | | | | | 0.83 | 0.96 |
| | n°3 | | | | | | | | | | | | 1.25 | 0.94 |
| | n°4 | | | | | | | | | | | | 0.55 | 0.93 |
| **Cochlea** | n°1 | 1 | 1 | 5 | 4 | 1 | 0.1 | 5 | 0.7 | 0.1 | 10 | 0.8 | 1.08 | 1.00 |
| **Culture** | n°1 | 23 | 16 | 5 | 4 | 1 | 0.9 | 5 | 0.9 | 0.1 | 5 | 0.8 | 0.81 | 0.98 |
| **Organoid** | n°1 | 5 | 12 | 5 | 2 | 1 | 0.9 | 5 | 0.8 | 0.1 | 10 | 0.8 | 0.83 | 1.00 |
| | n°2 | | | | | | | | | | | | 1.13 | 0.88 |

**Table S1**: for each dataset, this table summarizes the manually adjusted control parameters of Zellige used to extract the surfaces, and the corresponding quantification of the result quality in term of RMSE and coverage.

#### 4-1) Sensitivity analysis on a synthetic image.

The phantom image analysed shown in **Figure 2** is composed of one wavy sinusoidal surface (surface S1), presenting a homogeneous signal over the entire surface, and two superimposed surfaces (surfaces S2 and S3) presenting a signal restricted to a Voronoi polygonal mesh. The two polygonal meshes differ by the mesh size and topography. Surface S2 is supported by a plane and has inclination but no curvature, while surface S3 is supported by a paraboloid whose axis is at the center of the field of view. The study of such phantom images was very useful to develop Zellige by allowing us to test and better understand how to extract multiple surfaces showing various characteristics (in terms of contrast, texture, topography, etc) from a 3D image using a single set of control parameters. One next asked if small variations of one parameter in this manually defined parameter set would prevent the extraction of some surfaces: how robust is Zellige to parameter variations?

##### *Sensitivity to parameters controlling surface pixel selection*

The results of the sensitivity analysis with respect to the surface pixel selection step, namely the two classification threshold parameters ($T_A$, $T_{otsu}$) and the clean-up parameters (minimum island size $S_{min}$ and smoothing radiuses $\sigma_{xy}$ and $\sigma_z$) are shown in **Figure S2A**. Variations of the parameters $T_A$, $T_{otsu}$, and $S_{min}$ in their respective working intervals ($0 \leq T_A$, $T_{otsu} \leq 30$ and $0 \leq S_{min} \leq 50$) give rise only to small variations of the RMSE and the coverage of the three reconstructed height-maps. The reconstructions remain of high quality (RMSE < 1 and coverage > 95%) for all tested values of these parameters. Regarding the smoothing parameters ($\sigma_{xy}$ and $\sigma_z$), we observe a notable sensitivity on the reconstruction quality (**Figure S2A**). In the absence of smoothing along *xy* ($\sigma_{xy} = 0$), the reconstructed surface S 2 and S3 appear rough (with RMSE values much larger than 1) despite a good coverage (close to 100%). Thus, a minimum of smoothing in *xy* appears necessary to reconstruct surfaces presenting a mesh. The surface reconstruction is nevertheless robust as it remains high for a wide range of $\sigma_{xy}$ values ($1 \leq \sigma_{xy} \leq 7$). For too large values of this parameter ($\sigma_{xy} > 7$), the coverage of surface S3 gradually decreases, which is expected as this surface is strongly inclined away from the center of the field of view. The surface reconstruction is also quite robust against variations of the smoothing radius along the z axis, remaining of high quality for $1 \leq \sigma_z \leq 4$. For higher values of $\sigma_z$, the surfaces tend to merge, which results in incomplete reconstructions (low coverage) and poor accuracy (high RMSE). Also, using too large values of $\sigma_z$ is inadequate for strongly inclined surfaces.

*Sensitivity to parameters controlling surface assembly*

The surface reconstruction also depends on parameters controlling the assembly of the orthogonal surface elements (OSE), namely the parameters ($T_{OSE1}$, $C_1$, $R_1$) and ($T_{OSE2}$, $C_2$, $R_2$) controlling the fraction of OSE sizes allowed for seeds and the matching constraints required between OSEs and surfaces, during the first and second rounds of surface assembly, respectively. The results of the sensitivity analysis against variations of these parameters is shown in **Figure S2B.** The reconstruction turns out to be very robust to variations of the OSE seed size parameters $T_{OSE1}$, $T_{OSE2}$ and the connectivity parameters $C_1$ and $C_2$, remaining of high quality except for very stringent values ($T_{OSE1}$ <0.2 and $C_1$, $C_2 \simeq 1$). The quality of the reconstruction appears more sensitive to the parameters controlling the overlap constraint (parameters $R_1$ and $R_2$, see **Figure S2B**). Notably, surfaces that show regions of strong inclination tend to be more difficult to reconstruct using very stringent values (close to 1) of $R_1$ and $R_2$. This is expected since inclined surfaces typically display less overlap between OSEs. It is therefore important to keep this threshold value low enough to build surfaces with a steep slope.

In summary, the sensitivity analysis on this phantom image demonstrates the high efficiency and robustness of Zellige to extract multiple surfaces with disparate textures and contrasts, without requiring to know a priori the number or relative positions of the surfaces of interest.

#### 4-2) Sensitivity analysis on 3D images of biological samples

*Example 1: pupal fly specimen*

To examine the robustness of the segmentation by Zellige on biological images, we carried out a sensitivity analysis as described above for the phantom image (**Figure S3**). The sensitivity analysis using this fly specimen is described in details in the main text to emphasize that Zellige can robustly extract multiple surfaces from a complex biological image.

*Example 2: cochlea specimen*

Variations in the parameter $S_{\mathrm{min}}$ over its entire working interval ($0 \leq S_{\mathrm{min}} \leq 50$) result in stable high-quality values of the RMSE (< 1) and the coverage (> 95%) (**Figure S4A**), indicating that this parameter is not essential for this data set. The RMSE and coverage also remain rather stable upon variations of the thresholds ($T_{\mathrm{A}}$, $T_{\mathrm{otsu}}$) and smoothing parameters ($\sigma_{xy}$, $\sigma_z$), although the intervals ensuring high quality of the reconstructed surface are more restricted than for the fly sample: $0 \leq T_{\mathrm{A}} \leq 7$, $0 \leq T_{\mathrm{otsu}} \leq 8$, $\sigma_{xy} \geq 2$, and $0 \leq \sigma_z \leq 2$. An Otsu threshold that is too high leads to the exclusion of the area of interest in this example, because this region is of low signal intensity (**Figure S4A**). As this surface is supported by an epithelial mesh, an *xy* smoothing is necessary for accurate reconstruction. In addition, to avoid the fusion of the epithelial surface with the underlying structures forming the mesenchyme, the value of the parameter $\sigma_z$ should be chosen quite low. Regarding the surface assembly control parameters, the values of the RMSE and coverage of the reconstruction remain of high quality over their entire intervals of variation, except for extreme values of the connectivity parameters ($C_1$, $C_2 \simeq 1$) which lead to a drop in the overlap values. Thus (unlike the other surface extraction software PreMosa, FastSME and LocalZProjector), Zellige not only makes it possible to segment multiple surfaces, but also permits a surface characterized by low signal intensity and low contrast to be extracted selectively against surrounding structures of potentially higher contrast.

*Example 3: epithelial cell culture specimen*

For this image containing a single surface of interest of high roughness, the quality of the reconstructed surface is largely insensitive to variations of the $S_{min}$ parameter, remaining high over the whole interval $0 \leq S_{\mathrm{min}} \leq 50$. It may be noted, however, that larger values of $S_{min}$ result in lower RMSE and higher coverage values (**Figure S5A**) (for $S_{\mathrm{min}} = 50$, RMSE < 1 and the coverage is > 99%). This is not surprising as this specimen harbors out-of-plane dense spurious signal that becomes eliminated by the Island Search filter. The RMSE and coverage remain rather stable upon variations of the classification thresholds ($T_{\mathrm{A}}$, $T_{\mathrm{otsu}}$) and the smoothing parameters $\sigma_z$ (RMSE <1 and coverage > 99%) (**Figure S5A**). Yet, the reconstruction is more sensitive to variations of the smoothing parameter $\sigma_{xy}$, which gives best results for $\sigma_{xy} = 6$ (RMSE < 1 and coverage > 99%), while still giving good results around this value (RMSE < 1, coverage > 90%). With a few exceptions, the values of the RMSE and coverage are stable to variations of most parameters controlling the reconstruction of surfaces ($T_{\mathrm{OSE1}}$, $T_{\mathrm{OSE2}}$, $C_1$ and $C_2$), except for extreme values of the connectivity rates ($C_1$ or $C_2 \geq 0.95$) (**Figure S5B**). However, the reconstruction does show sensitivity to variations of the overlap parameters $R_1$ and $R_2$ that should be kept <30% for best results. This observation is likely explained by the particularly rough texture of the epithelium that leads to matching constraints more difficult to satisfy between OSEs and the reconstructed surface. Unlike other Premosa and FastSME, the approach developed with Zellige also makes it possible to extract rough surfaces with a good robustness. For this particular example, Zellige (through a manual adjustment of the parameters of steps 1 and 2) gives similar results to LocalZprojector (obtained as the best results in a parameter sweep as described in Hebert *et al*. 2021 [2].

*Example 4: inner ear organoid specimen*

Similarly to the above analyzed specimens, the RMSE and coverage remain stable upon variations of the classification thresholds ($T_A$, $T_{otsu}$) (RMSE ~1 and coverage > 80%) (**Figure S6A**). The reconstruction is more sensitive to the smoothing parameter $\sigma_{xy}$ and $\sigma_z$, which is expected due to the proximity of the two surfaces as well as the presence of regions of high inclination at their boundaries. Indeed, higher smoothing values will tend to blend the two surfaces and prevent their reconstruction: for $\sigma_z$ >3, the surface S1 is not reconstructed (null coverage) and another, chimeric surface is reconstructed instead, which is truly a blend of surfaces S1 and S2, as shown by the RMSE > 2. With a few exceptions, the values of the RMSE and coverage are stable to variations of most parameters controlling the reconstruction of surfaces ($T_{OSE1}$, $T_{OSE2}$, $C_1$ and $C_2$), except for extreme values of the connectivity rates ($C_1$ or $C_2$ ≥ 0.95). However, the reconstruction shows sensitivity to variations of the overlap parameters $R_1$ and $R_2$ that should be kept <30% for best results (**Figure S6B**). Again, this is expected considering the sharp inclination of the surfaces to be reconstructed at their boundary (where the surfaces touch the substrate). Nonetheless, Zellige makes it possible to robustly reconstruct two superimposed mesh-like surfaces including regions of strong inclination, using the same set of parameters.

### Supplemental note 5: Computational/processing time of Zellige

#### Computation time comparison between the various images tested

The core algorithms implementing the two main computational steps of Zellige (surface pixel selection step and surface assembly step) have linear computational complexities in the image size (**Figure S7**), i.e. both steps require a computation time that grows at most linearly in the number of pixels in the original 3D image. For step 1, this linear complexity results formally from the fact that the surface pixel selection amounts to a fixed number of tests applied sequentially to each pixel in the image. For step 2, linear complexity is less clear at the formal level. However, from a practical point of view this is a consequence of the fact that the number of constructed OSEs is bounded by the number of image pixels (in practice it is much smaller), and that the number of surfaces of interest to reconstruct is typically small. As shown below, in the examples we tested, the surface assembly step turns out to run about an order of magnitude faster than the surface pixel selection step.

**Figure S7** shows the execution times for running the surface extraction on the images considered in this study ranged between a few seconds to a few tens of seconds on a standard PC workstation (processor Intel Core i9 2,4 GHz). The computational load of Zellige thus allows the processing of large datasets in reasonable time. For all the images tested, the surface pixel selection step (step 1) is the most computationally costly, taking over >70% of the total execution time. The surface assembly step (step 2) usually runs in a few seconds, and rarely in more than 10s. This step 2 tends to be more computationally demanding for images containing numerous spurious structures surrounding the surface of interest, or when reconstructing a surface having a rough topography (as in the case of the cochlea and culture examples of **Figures 4** and **5**). The computational time of the surface assembly step turns out to be also sensitive to the parameters $T_{\mathrm{OSE1}}$ and $T_{\mathrm{OSE2}}$ setting the fraction of OSE sizes allowed for the OSE seeds during the two rounds of surface assembly. For values of 60% or more (when a majority of available OSE sizes are allowed as seeds), the computation time of step 2 raises significantly, occasionally reaching times comparable or larger than those of step 1. This is not unexpected since adopting large values for $T_{\mathrm{OSE1}}$ and $T_{\mathrm{OSE2}}$ means that large sets of OSE seeds have to be incorporated into surfaces before the construction is complete. It is thus advantageous to limit these parameters to values < 50%, which does not affect the quality of surface reconstruction. With this proviso, the most limiting step of Zellige regarding computational time is therefore the surface pixel selection step, which includes the two binary classifications schemes applied (amplitude and Otsu classifications), as well as the morphological clean-up operations embodied in the small island removal and the final gaussian interpolation. Note that the surface selection step essentially consists of a series filtering operations and of sequential pixel testing or computations, all of which could be to a large extent parallelized. The implementation of this step on a parallel platform (e.g. GPU or multi-core processors) might boost the computational time of Zellige sufficiently for real-time applications.

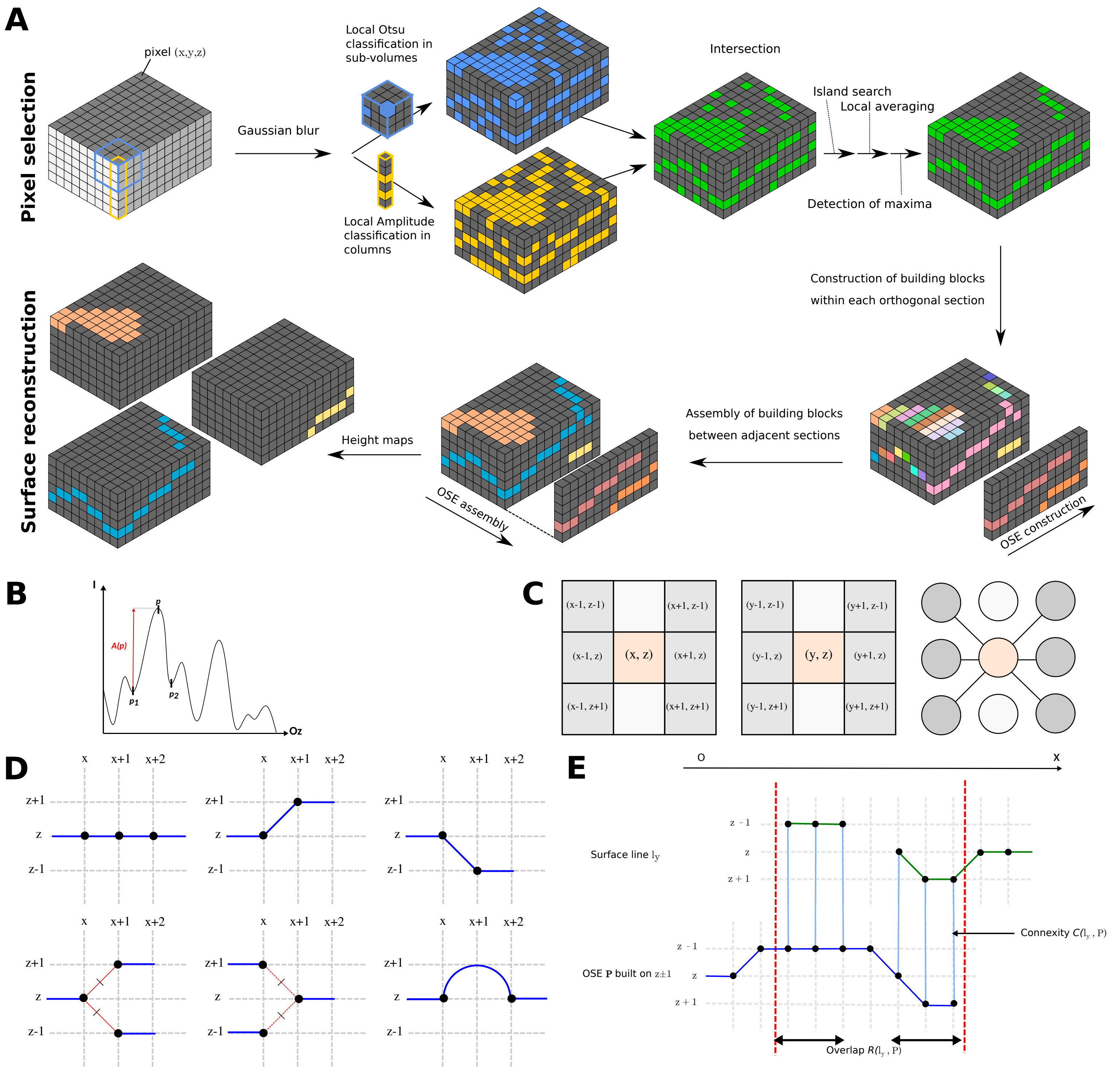

**Figure S1. Zellige implementation. (A)** The two main algorithmic steps of Zellige. Upper part: Surface pixel selection step (step 1). Lower part: Surface assembly step (step 2). **(B)** Determination of the amplitude *A(p)* of some local maximum *p* along the *z* axis, relative to its two closest local minima $p_1$ and $p_2$. **(C,D)** Connectivity rules used to connect putative surface pixels together for the construction of orthogonal surface elements (OSEs). These rules are based on a 6-connectivity relationship defined within each orthogonal (*xz* or *yz*) section as shown in (C). This relationship is extended to allow the presence of single straight gaps, while being constrained to forbid the occurrence of forking points. The allowed local neighborhood configurations along an OSE are illustrated in (D) in the case of a construction within *xz* sections. The OSEs are formally defined as the connected components of the graph $G_{OSE}$ defined by these rules within each orthogonal section. **(E)** Compatibility rules used in the surface assembly step, illustrated here in the case where OSEs have been constructed within *xz* sections, and assembly proceeds along the y axis. To validate the addition of a new OSE $\sigma$ constructed within section *y+1* (in blue) to the surface *S* under construction, whose intersection with section y is shown (surface line $l_y$, in green), two quantities are computed: the overlap $R(S,\sigma)$ is defined as the number of pixels of $\sigma$ that share the *x* coordinate of some pixel of $l_y$. The connectivity $C(S,\sigma)$ is defined as the fraction of the overlapping pixels of $\sigma$ that are 6-connected, in their respective *yz* section, to the corresponding point of $l_y$ (according to the 6-connectivity relationship shown in C, middle panel). In the depicted example, $R(S,\sigma)=6$ and $C(S,\sigma)=6/6=1$. The OSE will be added to *S* if $R(S,\sigma) \geq R_0$ and $C(S,\sigma) \geq C_0$, where $R_0$ and $C_0$ are tunable thresholds that set the stringency of the matching condition.

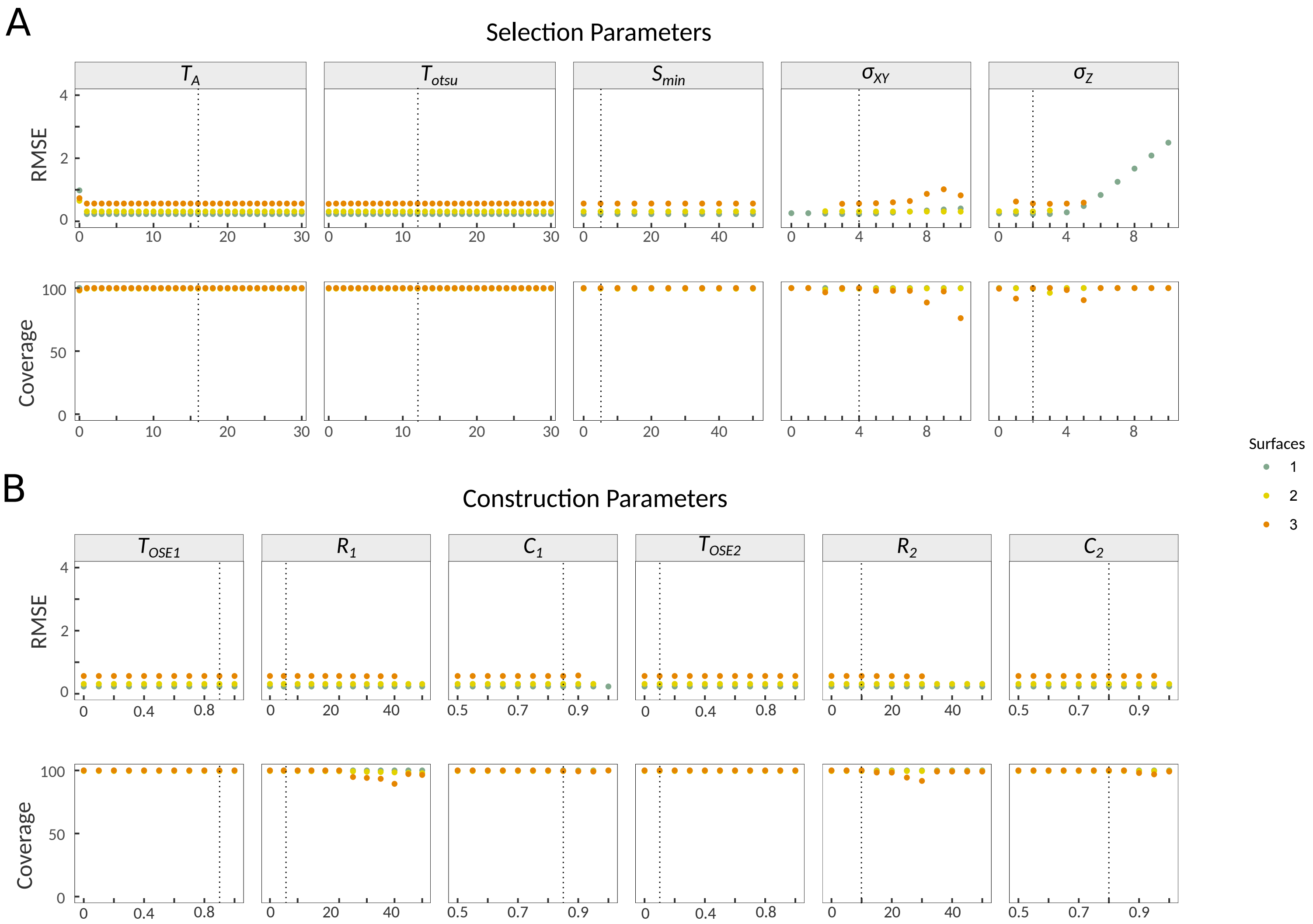

**Figure S2. Sensitivity analysis of Zellige on the phantom image. (A)** Surface pixel selection parameters. **(B)** Surface assembly parameters. Reference values are indicated by the dashed line.

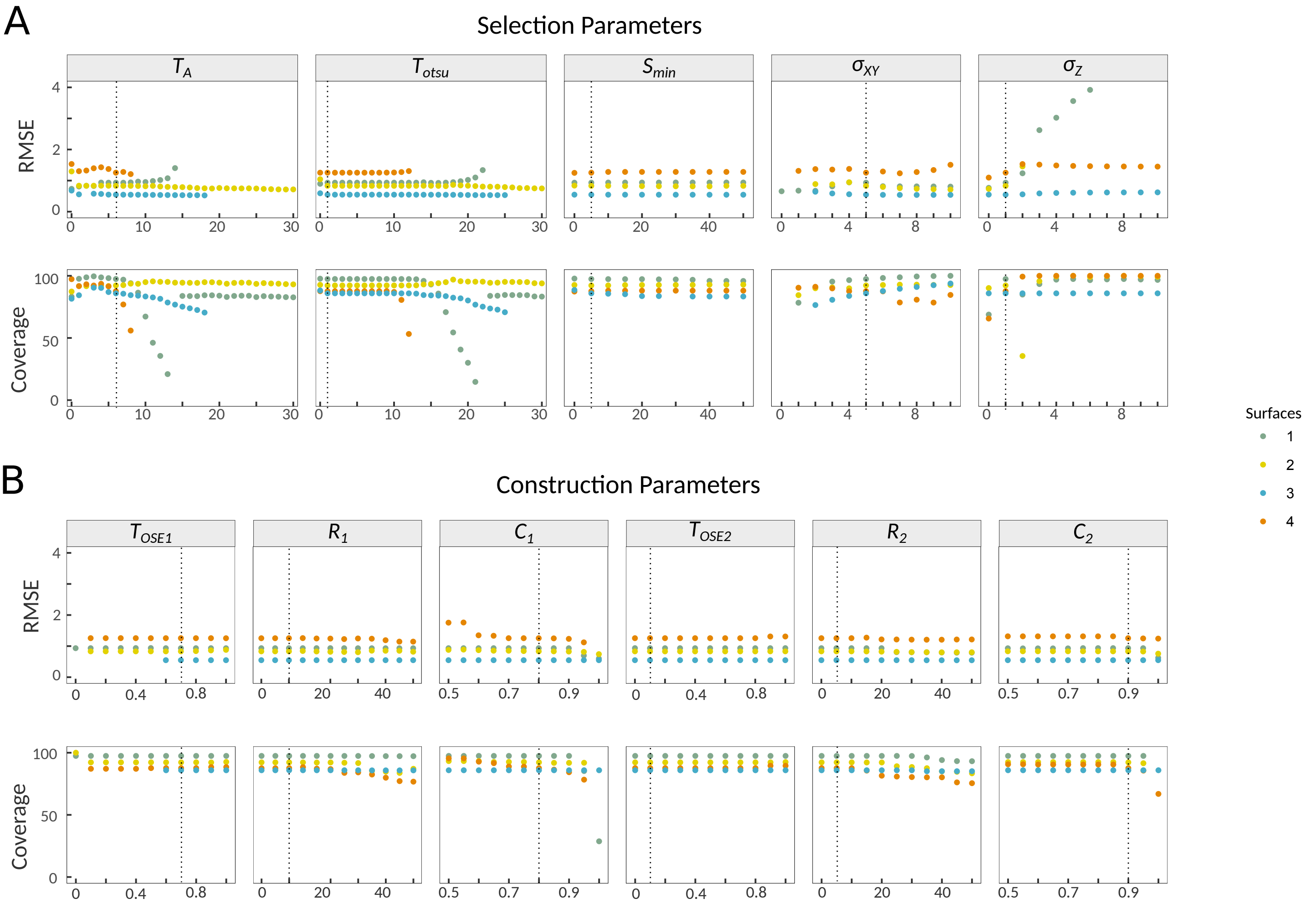

**Figure S3. Sensitivity analysis of Zellige on the pupal fly image. (A)** Surface pixel selection parameters. **(B)** Surface assembly parameters. Reference values are indicated by the dashed line.

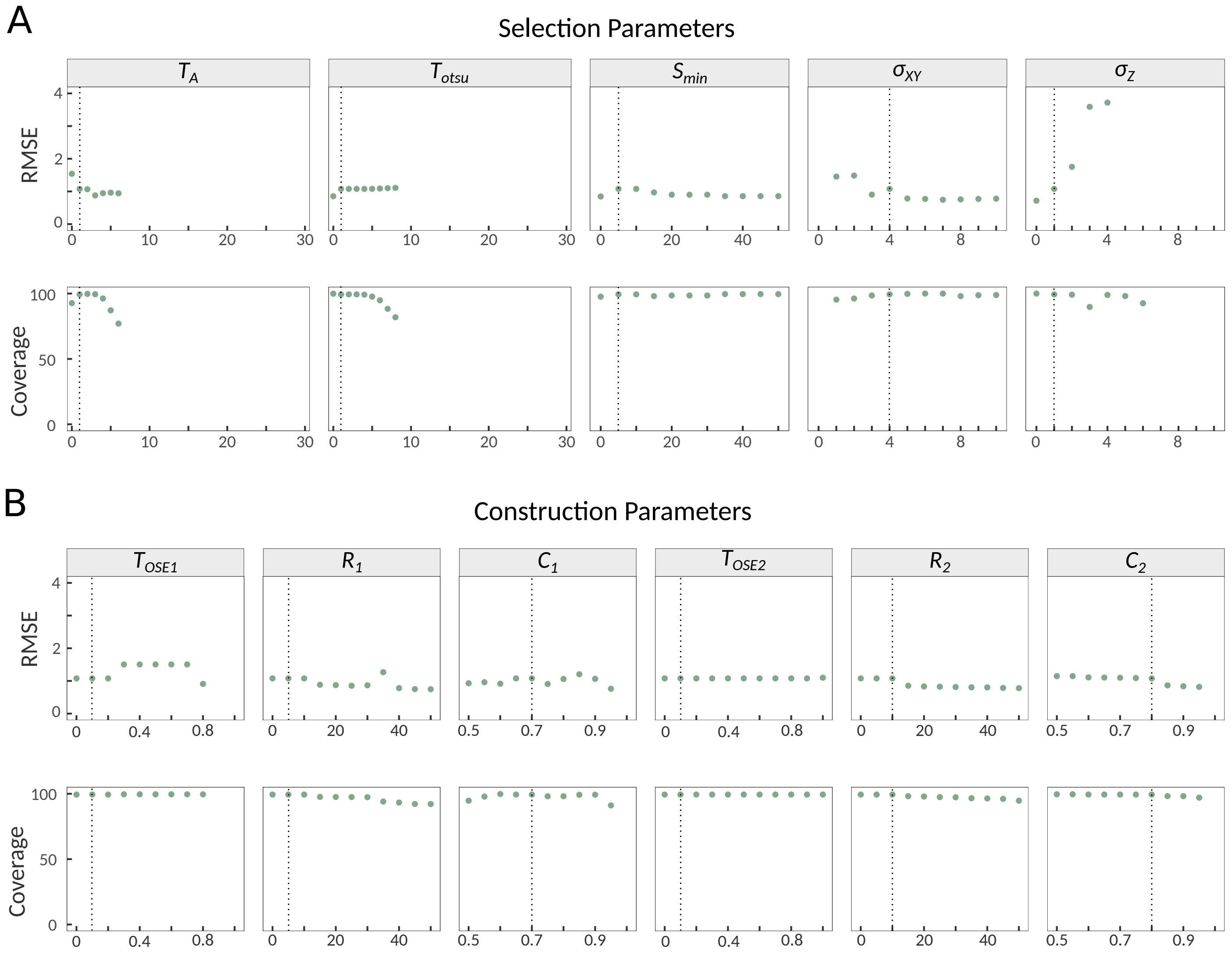

**Figure S4. Sensitivity analysis of Zellige on the cochlear epithelium image. (A)** Surface pixel selection parameters. **(B)** Surface assembly parameters. Reference values are indicated by the dashed line.

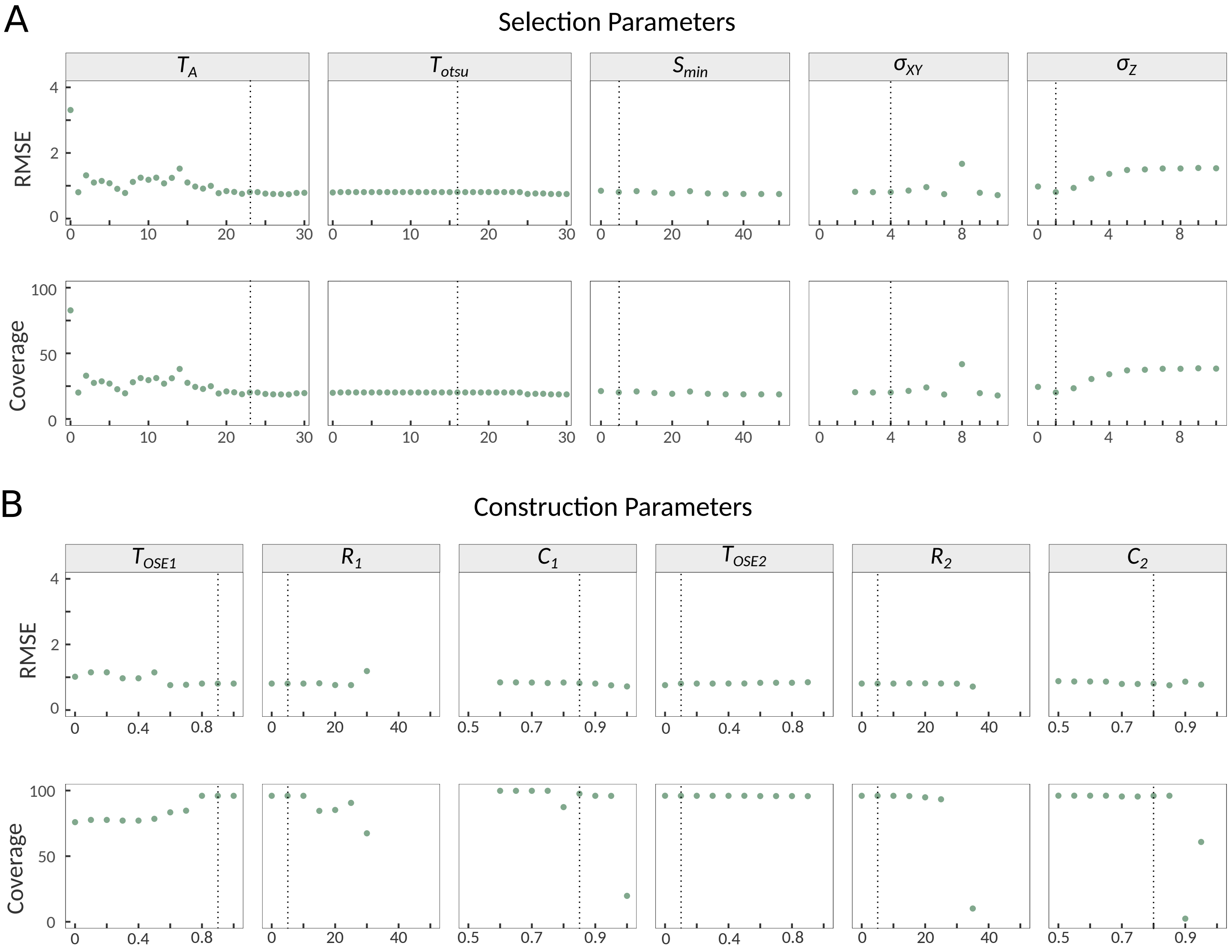

**Figure S5. Sensitivity analysis of Zellige on the primary epithelium culture image. (A)** Surface pixel selection parameters. **(B)** Surface assembly parameters. Reference values are indicated by the dashed line.

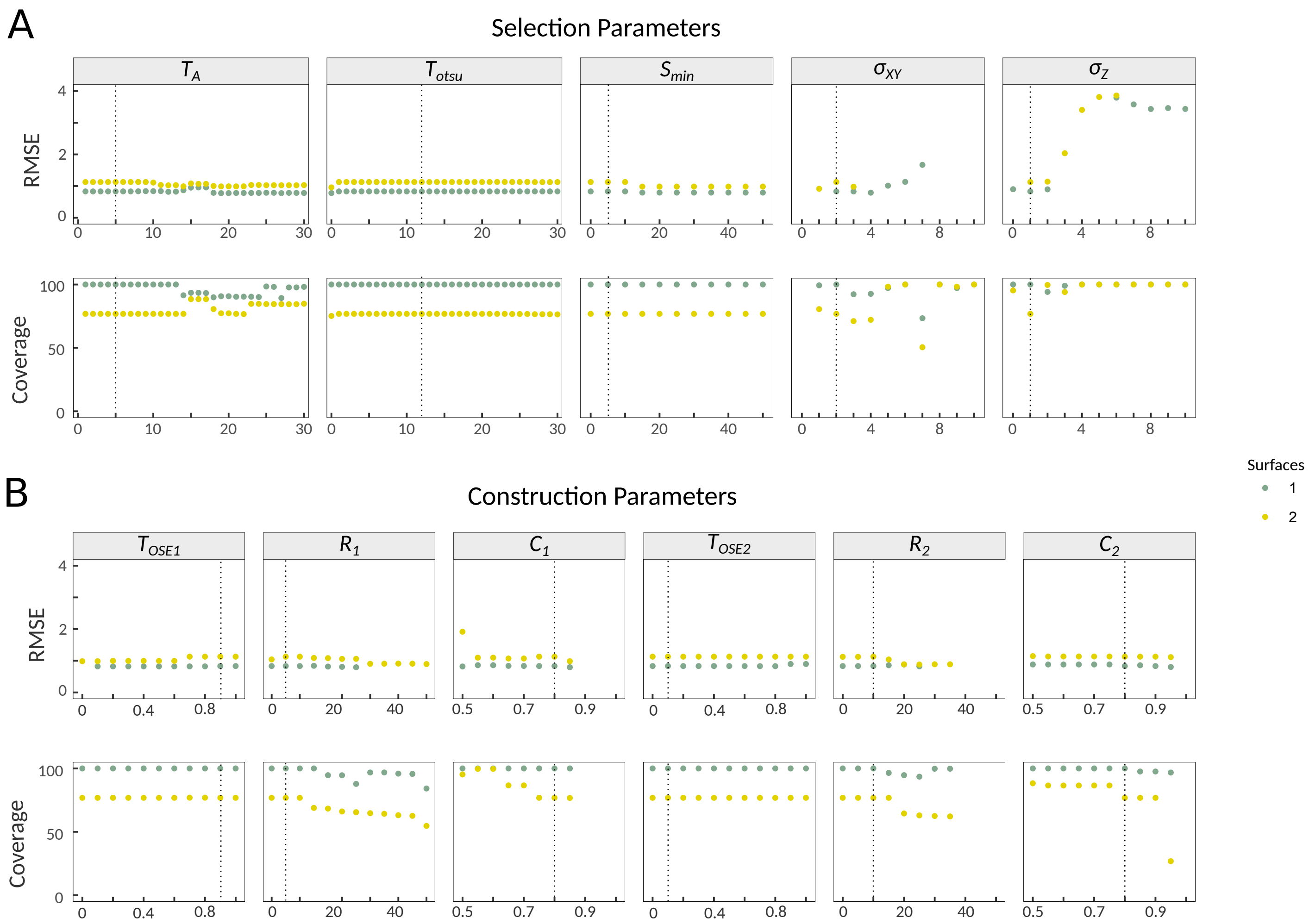

**Figure S6. Sensitivity analysis of Zellige on the inner ear organoid image. (A)** Surface pixel selection parameters. **(B)** Surface assembly parameters. Reference values are indicated by the dashed line.

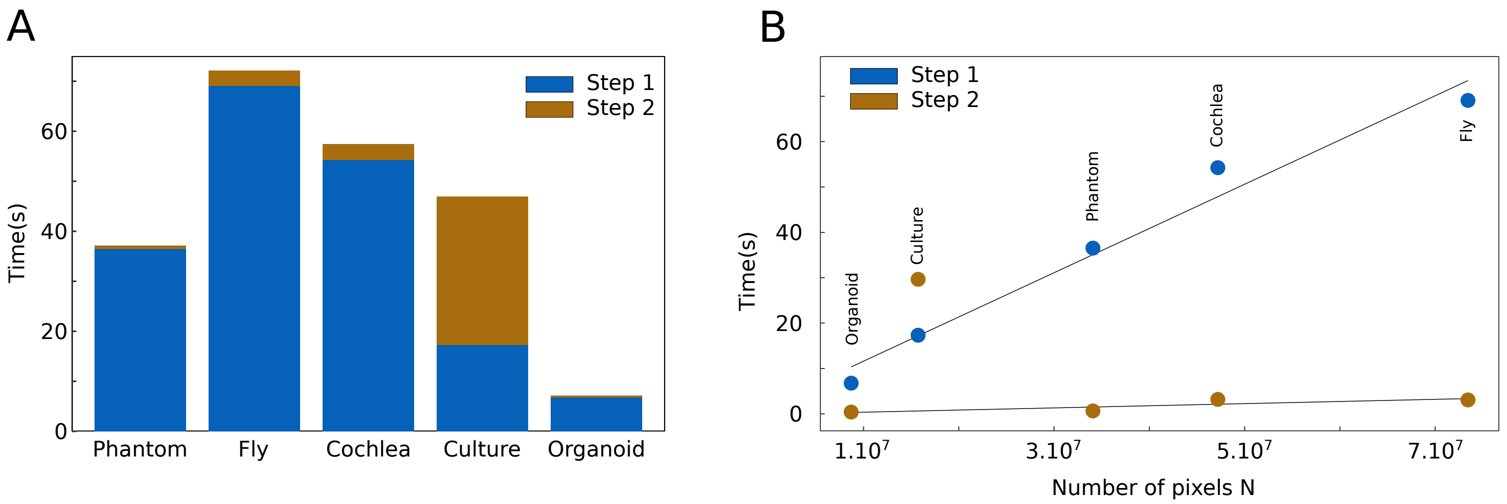

**Figure S7. Computational time analysis of Zellige. (A)** Computation times (step 1 in blue, step 2 in brown) for processing the various images tested on a PC notebook computer with (processor Intel Core i9 2,4 GHz with 32 Gb of Ram). Apart for the case of a highly rough surface (culture specimen) the computation time is largely dominated by the surface pixel selection step (step 1). **(B)** The same computation times are re-plotted as a function of image size *N* (number of pixels). Note the linear growth of the computational time of step 1 as a function of *N*, while that of step 2 shows much slower growth.

**FIGURE S1**

(A) The two main algorithmic steps of Zellige. Upper part: Surface pixel selection step (step 1). Lower part: Surface assembly step (step 2).

(B) Determination of the amplitude $A(p)$ of some local maximum $p$ along the $z$ axis, relative to its two closest local minima $p_1$ and $p_2$.

(C,D) Connectivity rules used to connect putative surface pixels together for the construction of orthogonal surface elements (OSEs). These rules are based on a 6-connectivity relationship defined within each orthogonal (*xz* or *yz*) section as shown in (C). This relationship is extended to allow the presence of single straight gaps, while being constrained to forbid the occurrence of forking points. The allowed local neighborhood configurations along an OSE are illustrated in (D) in the case of a construction within xz sections. The OSEs are formally defined as the connected components of the graph $G_{OSE}$ defined by these rules within each orthogonal section.

(E) Compatibility rules used in the surface assembly step, illustrated here in the case where OSEs have been constructed within *xz* sections, and assembly proceeds along the *y* axis. To validate the addition of a new OSE $\sigma$ constructed within section y+1 (in blue) to the surface $S$ under construction, whose intersection with section y is shown (surface line $l_y$, in green), two quantities are computed: the *overlap* $R(S,\sigma)$ is defined as the number of pixels of $\sigma$ that share the $x$ coordinate of some pixel of $l_y$. The *connectivity* $C(S,\sigma)$ is defined as the fraction of the overlapping pixels of $\sigma$ that are 6-connected, in their respective yz section, to the corresponding point of $l_y$ (according to the 6-connectivity relationship shown in C, middle panel). In the depicted example, $R(S,\sigma)$=6 and $C(S,\sigma)$=6/6=1. The OSE will be added to $S$ if $R(S,\sigma) \geq R_0$ and $C(S,\sigma) \geq C_0$, where $R_0$ and $C_0$ are tunable thresholds that set the stringency of the matching condition.

**FIGURE S2**

Sensitivity analysis of Zellige on the phantom image. (A) Surface pixel selection parameters. (B) Surface assembly parameters. Reference values are indicated by the dashed line.

**FIGURE S3**

Sensitivity analysis of Zellige on the pupal fly image. (A) Surface pixel selection parameters. (B) Surface assembly parameters. Reference values are indicated by the dashed line.

**FIGURE S4**

Sensitivity analysis of Zellige on the cochlear epithelium image. (A) Surface pixel selection parameters. (B) Surface assembly parameters. Reference values are indicated by the dashed line.

**FIGURE S5**

Sensitivity analysis of Zellige on the primary epithelium culture image. (A) Surface pixel selection parameters. (B) Surface assembly parameters. Reference values are indicated by the dashed line.

**FIGURE S6**

Sensitivity analysis of Zellige on the inner ear organoid image. (A) Surface pixel selection parameters. (B) Surface assembly parameters. Reference values are indicated by the dashed line.

**FIGURE S7**

Computational time analysis of Zellige. (A) Computation times (step 1 in blue, step 2 in brown) for processing the various images tested on a PC notebook computer with (processor Intel Core i9 2,4 GHz with 32 Gb of Ram). Apart for the case of a highly rough surface (culture specimen) the computation time is largely dominated by the surface pixel selection step (step 1). (B) The same computation times are re-plotted as a function of image size *N* (number of pixels). Note the linear growth of the computational time of step 1 as a function of *N*, while that of step 2 shows much slower growth.
